# Pericyte-tumor crosstalk facilitates metastatic tumor cell latency through PIEZO1-activated lysophospholipid transfer

**DOI:** 10.1101/2025.02.15.638347

**Authors:** Tamara McErlain, Morgan J Glass, Elizabeth C McCulla, Caitlin A Madden, Lauren E Ziemer, Kirsten E Overdahl, Alan K Jarmusch, Michael J Kruhlak, Alexander T Chesler, Howard H Yang, Maxwell P Lee, Cristina M Branco, Meera Murgai

## Abstract

Tumor dissemination is increasingly recognized to begin early in tumor development. Although most of these early disseminated cells are cleared, some survive and persist below clinical detection, acting as reservoirs for metastatic relapse. Metastatic tumor cells often rely on interactions with local stromal cells to support their colonization. In this study, we propose that pericyte-tumor cell interactions promote dormancy induction in the early metastatic lung, enhancing disseminated tumor cell (DTC) persistence. Extravital imaging demonstrated that DTCs interact with pericytes upon extravasation into the lung. Co-culture experiments were used to assess DTC fate after pericyte contact and revealed that transient contact with pericytes reduced the proliferation of metastatic 4T1 breast cancer cells but had no effect on non-metastatic 67NR cells. *In vivo*, transient pericyte contact resulted in higher lung metastatic burden, driven by small, non-proliferative lesions (<6 cells), 10 days after intracardiac injection. These lesions exhibited reduced KI67 staining and EdU incorporation compared to those from monocultured cells. We further observed that primary lung pericytes transferred lyso-phospholipids (lyso-PLs) specifically to metastatic 4T1 cells through direct contact. Gene expression analysis indicated that transient pericyte contact activated pathways related to syncytium formation in metastatic cells. In normal physiology, pericytes act in a syncytium to regulate blood flow via mechanosensitive channels in response to blood pressure changes. We hypothesize that tumor cells exploit these mechanosensitive responses to trigger lyso-PL transfer from pericytes. Supporting this, calcium imaging showed higher calcium activity in pericytes co-cultured with 4T1 cells, and calcium channel inhibitors significantly reduced lyso-PL transfer. Pharmacological activation of pericyte calcium channels induced lyso-PL release, which was subsequently taken up by tumor cells. Conditioned medium from activated pericytes, containing free lyso-PLs, recapitulated the reduced proliferation observed in transient co-culture. Finally, we found our pericyte-induced dormancy signature to be associated with tumor dormancy and distant metastasis free survival latency in breast cancer patients. Together, these findings suggest that early DTCs may exploit pericyte signaling mechanisms to enter dormancy, facilitating their persistence at metastatic sites and contributing to future relapse.

## Introduction

Metastasis is responsible for most cancer-related deaths^1^. Notably, metastatic relapse is observed in a significant number of patients previously considered disease free and can occur months to decades after primary diagnosis and treatment^2,3^. A growing body of evidence suggests that metastatic dissemination occurs early during tumor progression^4–7^, with both preclinical models and clinical analyses revealing that disseminated tumor cells (DTC) can persist in a state of prolonged quiescence within metastatic niches^8–10^. DTC capable of surviving in a reversible cell cycle arrest, referred to as dormant, are thought to be responsible for metastatic relapse^2^.

The fate of DTC is partly dictated by the microenvironment encountered upon arrival to metastatic niches^11–14^, which can influence if, and when, tumor cells remain dormant or proliferate to form metastatic lesions. Much of the field has focused on the metastatic immune stroma which influence whether tumor cells persist and proliferate through immune-surveillance and - suppression activities^15–19^. Less is known, however, about the contributions of non-immune stromal cells on metastatic outcome. We and others have previously identified NG2-expressing pericytes as critical mediators of premetastatic microenvironments^11,20^. Pericytes play an early role in remodeling the microenvironment to facilitate tumor cell arrival and growth^11^. Additionally, the perivascular niche is recognized as critical for successful metastatic colonization^14,21,22^, with pericytes being one of the first cells encountered by tumor cells on arrival in a distant site. Here, we investigated the hypothesis that pericytes initiate critical DTC fate decisions upon arrival to potential metastatic niches. Direct contact between pericytes and tumor cells results in mechanosensitive calcium channel activation, resulting in rapid transfer of pericyte-derived lyso-PLs to metastatic tumor cells, promoting tumor cell latency. These data uncover a novel function for pericyte-dependent metastatic spread and suggest that pericyte-tumor intercellular communication is critical for determining DTC fate decisions.

These results align with previous observations of dormant DTC residing within perivascular niches in the lung, bone and brain^21,22^, as well as the dormancy-promoting microenvironment established by endothelial cells^14,21^. Our study extends this understanding by identifying perivascular pericytes as key contributors to the dormancy-inducing effects of the vascular niche. Given the lack of metastasis-specific targeted therapies and the clinical challenge of metastatic recurrence, the proposed mechanism by which pericytes shift the fate of DTCs towards dormancy may offer a novel strategy to limit metastatic relapse.

## Results

### Transient pericyte-tumor engagement initiates latency in disseminated tumor cells

To characterize pericyte-tumor interactions upon DTC arrival to the lung, we first assessed tumor cell engagement with pericytes during extravasation. We developed NG2-ERT^2^-cre Rosa-STOP-flox-zsGreen (NG2^zsGreen^) mice to reliably trace pericyte behaviors *ex vivo*, where all NG2-positive pericytes express the zsGreen fluorescent protein. Cardiac injection of mCherry-expressing metastatic 4T1 breast cancer cells into the NG2^zsGreen^ model revealed that 49% of tumor cells encounter lung pericytes during extravasation over the course of 2 hours of live imaging **(Fig 1A-C; Supplemental Fig 1A)**. The proportion of pericyte-tumor interactions reached saturation at 20 minutes post-tumor injection, after which the percent engagement with pericytes remained relatively unchanged even as the total number of extravasating tumor cells continued to increase **(Fig 1C)**. We have previously demonstrated that chronic, long-term exposure to tumor-derived factors promotes perivascular cell proliferation which in turn enhances tumor proliferation^11^. Our current *ex vivo* observations, however, suggest that understanding the impact of transient pericyte-tumor cell crosstalk is also relevant to metastatic colonization.

**Figure 1.**
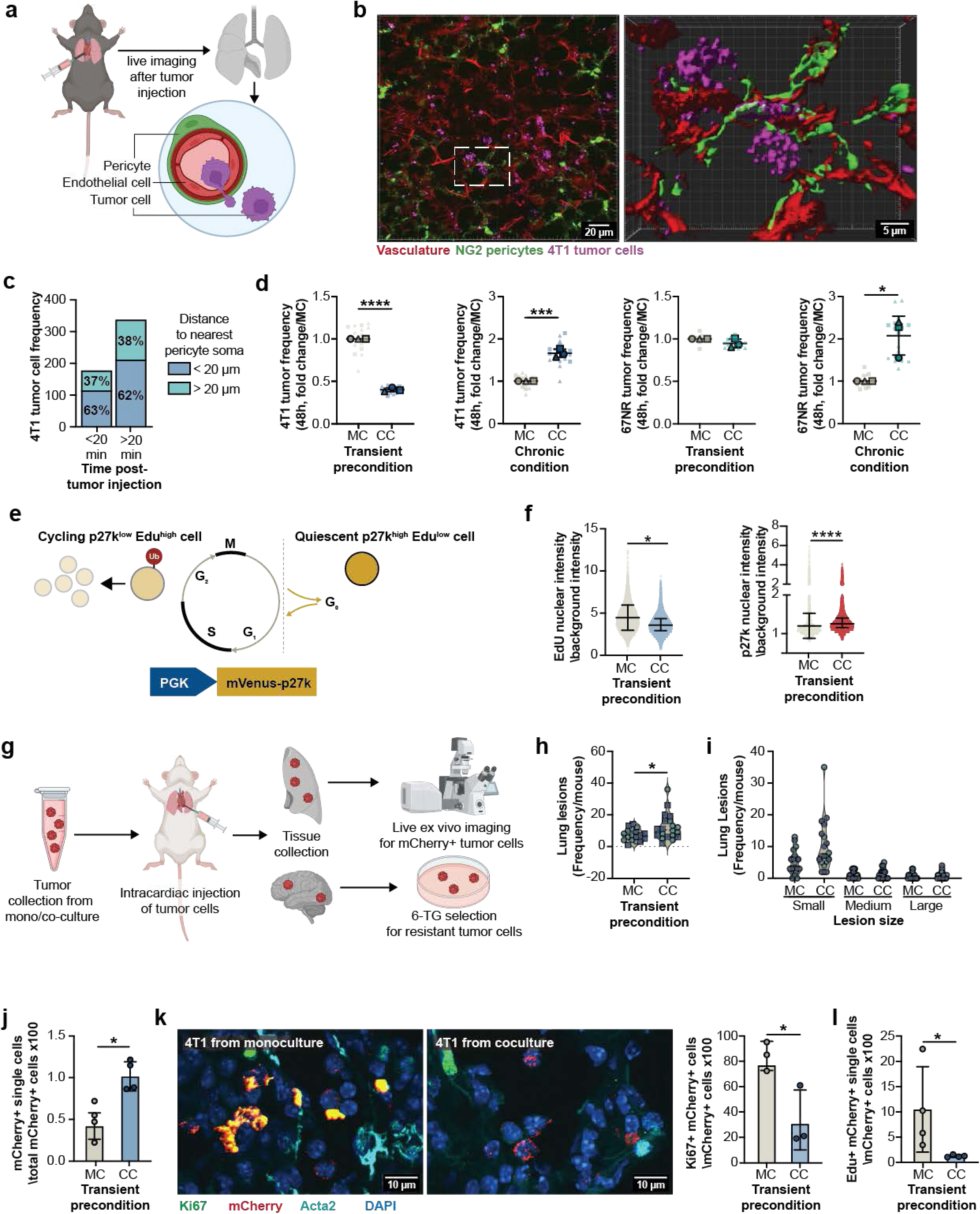
Transient pericyte engagement promotes disseminated tumor cell latency. **(a)** Experimental workflow: NG2-zsGreen pericyte lineage tracing mice were intracardiac injected with mCherry expressing 4T1 cells and a vascular tracer upon euthanasia. Lungs were dissected and imaged immediately using confocal microscopy. **(b)** Confocal images demonstrate NG2+ pericytes (green) closely associated with the lung vasculature (red) and in direct contact with 4T1 tumor cells (magenta). Surface rendering with Imaris software highlights 4T1-pericyte interactions. **(c)** Frequency of 4T1 tumor cells < or > than 20 μm from the nearest pericyte soma at < or > 20 min post tumor cell injection (3 lungs analyzed). **(d)** Tumor cell proliferation 48 h after isolation from transient co-culture with pericytes (termed transient pre-condition) or maintained in co-culture with pericytes (termed chronic condition); compared to tumor cell monoculture controls. Data are displayed as Superplots (Large symbols: means of experimental replicates; Small symbols: technical replicates for each experimental replicate; Lines: mean ±SD of replicate means). Statistical analysis was performed on the mean experimental replicate data only (Unpaired t test, ****p<0.0001, ***p<0.001). **(e)** Reporting quiescent cells: p27k mVenus expression is higher in quiescent cells than cycling cells, p27k high cells should also be EdU low. **(f)** 4T1 nuclear EdU intensity and p27k intensity 48 h after transient precondition with pericytes, compared to monoculture controls. (Kolmogorov-smirnov test, *p<0.05, ****p<0.0001). **(g)** Experimental workflow: 4T1 cells from transient pericyte preconditioning or monoculture control, were intracardially injected into mice. After 10 days, mice were euthanized, and lungs were collected for live *ex vivo* confocal imaging to identify mCherry-positive lesions on the lung surface followed by processing for immunofluorescence or organs were digested and cultured with the selection agent 6-thioguanine (6-TG) to isolate and quantify colony-forming tumor cells. **(h)** Total lung lesions per mouse from monoculture or transient co-culture groups. Statistical significance was assessed by unpaired t test (*p<0.05). Data is plotted per mouse and independent experiments can be differentiated by symbol shape and color (N=2; n=10/group/experiment). **(i)** Surface lung lesions were categorized by size into small (<6 cells), medium (<100 cells), and large (>100 cells). **(j)** Quantification of mCherry+ single cells in the lung presented as a proportion of total mCherry+ cells (One-way ANOVA, *p<0.05). **(k)** Representative images of lung sections from mice that received injection of tumor cells from transient preconditioning with pericytes or monoculture control, stained for Ki67, mCherry, Acta2 and DAPI. The percentage of ki67+ mCherry cells was quantified (Unpaired t test, *p<0.05). **(l)** Quantification of EdU+ mCherry+ single cells in the lung presented as a proportion of total mCherry+ cells (One-way ANOVA, *p<0.05).

To determine the impact of transient pericyte contact on metastatic tumor cell proliferation, we quantified tumor cell numbers after transient and chronic pericyte-tumor engagement *in vitro*. 4T1 tumor cells were added to pericyte-containing cultures for 20 minutes, after which the tumor cells were re-isolated, counted, and plated separately to assess tumor cell accumulation after 48 hours. Re-isolation of 4T1 tumor cells after transient co-culture resulted in a 99% pure tumor cell population **(Supplemental Fig 1B-C)**. Consistent with our previous observations using a vascular smooth muscle cell line^11^, chronic 48-hour co-culture with primary lung pericytes resulted in higher 4T1 tumor cell numbers compared to 4T1s grown in monoculture **(Fig 1D)**. Unexpectedly, we discovered that 4T1 cell numbers were significantly reduced after transient pericyte-tumor co-culture when compared to those grown in monoculture **(Fig 1D)**, which was recapitulated in an alternative metastatic tumor cell line, E0771 **(Supplemental Fig 1D)**. This opposing impact of transient and long-term pericyte co-culture on highly metastatic 4T1 tumor cells was not observed with non-metastatic 67NR cells which exhibited no change in cell number between transient co-culture and monoculture **(Fig 1D)**.

Reduced cell numbers after transient pericyte co-culture could be explained by either a reduction in proliferation, or a decrease in cellular fitness. To examine a role for altered cell cycle dynamics in tumor cells after transient pericyte co-culture, we measured the proportion of tumor cells that were either cycling or quiescent. We examined EdU incorporation over 16 hours in culture to identify cycling cells and utilized the p27k-mVenus reporter of quiescence to identify cells that persist in a long-term G0 phase of the cell cycle and exhibit high levels of p27k-mVenus^23^ **(Fig 1E, Supplemental Fig 1E)**. Transient pericyte co-culture resulted in a decrease in EdU incorporation and higher p27k-mVenus fluorescence intensity in 4T1 tumor cells that were recovered from transient pericyte co-culture when compared to monoculture, indicating that a shift towards quiescence may explain our observations **(Fig 1F)**.

To assess the cellular fitness of metastatic tumor cells after transient contact with pericytes, we injected tumor-naïve mice with mCherry-expressing 4T1 tumor cells that were either collected from transient pericyte co-culture, or non-pericyte containing monoculture **(Fig 1G)**. There was a greater total tumor burden in the lungs of mice that received transiently pericyte-educated 4T1 tumor cells at 10 days post-injection, implying that tumor cell survival was not impaired by transient pericyte co-culture **(Fig 1H)**. This greater tumor burden was entirely due to a higher count of persistent single cells, with no change in the frequency of larger, multicellular lesions **(Fig 1I-J)**. Similarly, we observed a greater number of mice with persistent 4T1 brain DTCs after transient pericyte co-culture compared to monoculture **(Supplemental Fig 1F)**. The observed increase in microlesions was unlikely to be attributed to an increase in extravasation efficiency, as no significant change was observed in the number of 4T1 tumor cells recovered nor identified in the lungs of mice 3 days post-injection that received pericyte-educated tumor cells compared to those that received pericyte-naïve tumor cells **(Supplemental Fig 1G-H)**.

Our *in vitro* observations reveal that transient pericyte contact results in a quiescent 4T1 subpopulation **(Fig 1F)**. To determine whether transient pericyte contact results in quiescent tumor cells *in vivo*, we examined the proliferative status of DTCs at 3- and 10-days post-injection. Proliferation marker Ki67 staining in mCherry+ tumor cells found in the lungs of mice that received tumor cells from pericyte co-culture was significantly lower than that seen in monocultured cells **(Fig 1K)**. Similarly, the number of EdU+ mCherry+ single tumor cells found in the lungs of mice that received pericyte co-cultured cells at 3 and 10 days after cardiac injection was also significantly lower **(Fig 1L, Supplemental Fig 1I)**. Together these data suggest that transient pericyte-tumor engagement promotes tumor cell latency both *in vitro* and *in vivo*.

### Metastatic tumor cells acquire pericyte-derived phospholipids through transient contact

To identify potential mechanisms of pericyte-directed tumor latency, we analyzed the gene expression pathway enrichment in 4T1 tumor cells after transient pericyte co-culture. Compared to monoculture, we observed a positive enrichment in cell viability and senescence pathways, and negative enrichment in necrosis and apoptosis pathways **(Fig 2A)**. These observations were consistent with our data suggesting that transient pericyte engagement enriches for 4T1 tumor cells with altered cell cycle dynamics **(Fig 1)**. Pathways associated with lipid metabolism were also enriched compared to monoculture **(Fig 2A, Supplemental Figure 2A)**. Lipid metabolism has previously been linked to cell survival and quiescence in many settings, including tumor dormancy and proliferative outbreak^24–29^. Circulating lipids also modulate perivascular cell phenotypic switching and vascular function in cardiovascular diseases^30–32^. To investigate the role of lipid metabolism in pericyte-directed tumor dormancy, we performed untargeted metabolomics screening in 4T1 and 67NR tumor cells. Our analysis revealed an enrichment in many phospholipids, including lysophosphatidylcholines (lyso-PCs) and lysophosphatidylethanolamines (lyso-PEs) that were unique to 4T1 cells from transient pericyte co-culture, and were not enriched in 4T1 cells from monoculture, nor 67NR cells from either co- or monoculture **(Fig 2B, Supplemental Fig 2B, Supplemental Table 1-2)**. Lyso-PCs and lyso-PEs play significant roles in cardiovascular function and cellular fate decisions, including apoptosis and tumor cell dormancy^33–37^. We therefore tested if phospholipid movement between pericytes and 4T1 cells is altered during transient crosstalk.

**Figure 2.**
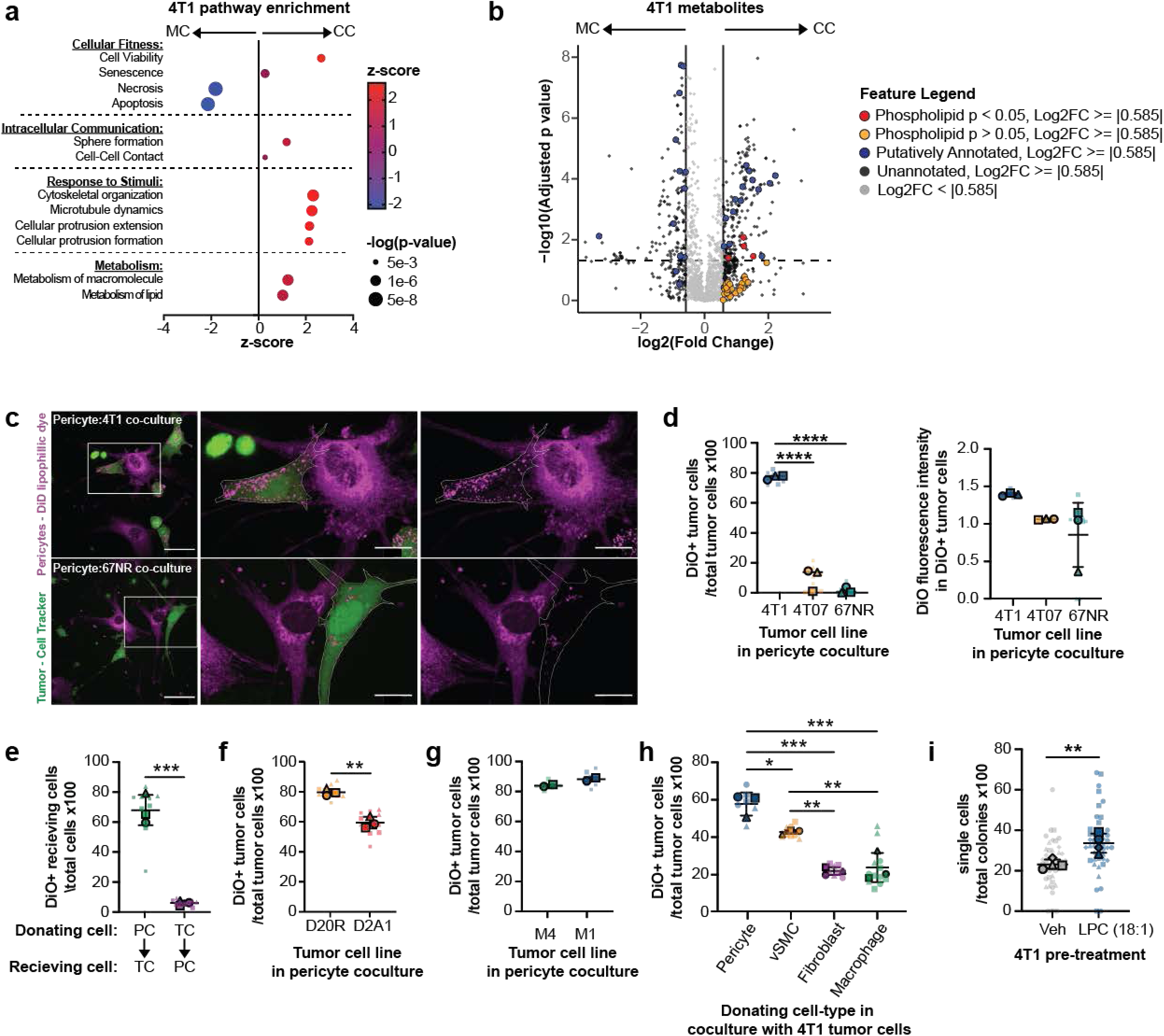
Metastatic tumor cells acquire phospholipids upon transient contact with pericytes. **(a)** Pathway enrichment in 4T1 tumor cells exposed to transient pericyte precondition or monoculture control. **(b)** Volcano plot of metabolite differences between 4T1 cells exposed to transient pericyte precondition or monoculture control. **(c)** Representative images of lipid transfer to 4T1 and 67NR tumor cells over 2 h. Pericytes labelled with DiD lipophilic dye (magenta) and tumor cells labelled with cell tracker (green). **(d)** Quantification of DiO+ tumor cells as a percentage of the total tumor cell population and DiO fluorescence intensity in the DiO+ subpopulation (One-way ANOVA with Tukey’s multiple comparisons test, ****p<0.0001). **(e)** Quantification of bidirectional transfer of phospholipids between pericytes and tumor cells (Unpaired t test, ***p<0.001). Percentage of DiO+ tumor cells in mouse cell lines **(f)** D20R and D2A1 and human cell lines **(g)** M1 and M4 (Unpaired t test, **p<0.01). **(h)** Quantification of efficiency of DiO transfer to tumor cells from various stromal population (One-way ANOVA with Tukey’s multiple comparisons test, *p<0.05, **p<0.01, ***p<0.001). **(i)** Percentage of single cell colonies in soft agar 7 days following transient treatment with Lyso-PC 18:1 (Unpaired t test, **p<0.01). Data are displayed as Superplots (Large symbols: means of experimental replicates; Small symbols: technical replicates for each experimental replicate; Lines: mean ±SD of replicate means). Statistical analysis was performed on the mean experimental replicate data only.

Live-cell imaging studies during transient pericyte-tumor co-culture, using fluorescent phospholipid analogs such as DiO, revealed that 4T1 tumor cells acquired pericyte-derived phospholipids **(Fig 2C-D)**. The reverse process of tumor-derived phospholipid uptake by pericytes was infrequent and occurred at minimal levels, suggesting a unidirectional mechanism of phospholipid transfer **(Fig 2E)**. High levels of phospholipid transfer were also seen in metastatic D2A1 cells, but not in low metastatic 4T07 and 67NR cells **(Fig 2D, F)**. Human tumor and epithelial cell lines, MCF10 Ca1a.c11 (M4) and MCF10A1 (M1), were also observed to receive pericyte-derived lipids during transient co-culture with a human lung pericyte cell line **(Fig 2G)**. Interestingly, the dormant-prone D20R cell line exhibited high levels of pericyte-derived phospholipid uptake, proposing that phospholipid dynamics may also be relevant in other established models of tumor dormancy **(Fig 2F, Supplemental Fig 2C)**^38–41^.

Other studies have noted that tumor cells may scavenge lipids and metabolites from their local environment, including those derived from fibroblasts and macrophages^13,42–45^. We therefore evaluated the efficiency of phospholipid transfer to 4T1 tumor cells during transient co-culture with a range of stromal cell-types. Pericyte co-culture yielded a significantly higher proportion of tumor cells with phospholipid uptake when compared to fibroblasts and macrophages **(Fig 2H)**. Interestingly, phospholipid transfer was also observed with vascular smooth muscle cells which share pericyte functions as perivascular support cells, indicating that similar mechanisms for phospholipid transfer may be employed by these two cell types.

Pericyte-tumor cellular content transfer during transient contact was specific to phospholipids, in that the transfer of other pericyte-derived cellular contents, including intracellular and membrane-targeted proteins, was very infrequent **(Supplemental Fig 2 D-G)**. Importantly, although we observed the uptake of pericyte-derived neutral lipids such as cholesterol to tumor cells, this transfer was not specific to highly metastatic 4T1 tumor cells compared to 67NR cells as was observed with phospholipids **(Supplemental Fig 2H)**. Furthermore, pericyte-derived cholesterol transfer required up to 24 hours to occur to detectable levels **(Supplemental Fig 2H)**. These observations were consistent with our mass spectrometry results that identified an enrichment of phospholipids after transient pericyte co-culture, but no enrichment in other potential metabolites **(Fig 2B, Supplemental Tables 1-2).** Using a candidate phospholipid identified by mass spectrometry, lyso-PC 18:1, we assessed if this agent was sufficient to recapitulate the observed persistence of single 4T1 tumor cells *in vivo* after transient co-culture with pericytes. Transient treatment of 4T1 tumor cells with lyso-PC 18:1 resulted in a greater number of single cells in a soft agar colony formation assay 7 days later **(Fig 2I)**.

Taken together these data suggest that the pericyte-tumor intracellular trafficking during transient engagement occurs through perivascular cell-specific mechanisms and with phospholipid-specific cargoes, to promote tumor cell quiescence, in contrast to what has previously been observed by fibroblasts and macrophages^42,45^.

### Metastatic tumor cells initiate phospholipid transfer through pericyte mechanosensitive channel activation

Recent studies have noted that tumor cells take up exogenous lipids from the surrounding microenvironment when they are secreted into the extracellular space by surrounding donor cells such as fibroblasts and macrophages^42,45^. We observed that lipid transfer by fibroblasts and macrophages was highly inefficient when compared to pericytes **(Fig 2)**. Nonetheless, a common mechanism of transfer via secretion into the extracellular microenvironment could explain our observations. We therefore tested whether phospholipid transfer required direct cell-cell contact. These cell culture configurations included culturing 4T1 tumor cells in pericyte-conditioned medium and pericyte-tumor co-culture across a transwell membrane that prevented direct cell contact **(Fig 3A)**. Surprisingly, only direct pericyte-tumor cell contact was sufficient to observe maximal phospholipid transfer to tumor cells during 2 hours of co-culture **(Fig 3B, Supplemental Fig 3A)**. We observed that only 1% of tumor cells received pericyte-derived phospholipids after exposure to pericyte conditioned medium, and levels of phospholipids transferred were also significantly lower than direct co-culture conditions **(Fig 3B, Supplemental Fig 3A)**. These data indicate that pericyte-derived phospholipid transfer utilizes a mechanism more efficient and requires direct contact, in contrast to what has been described for fibroblasts and macrophages^42,45^.

**Figure 3.**
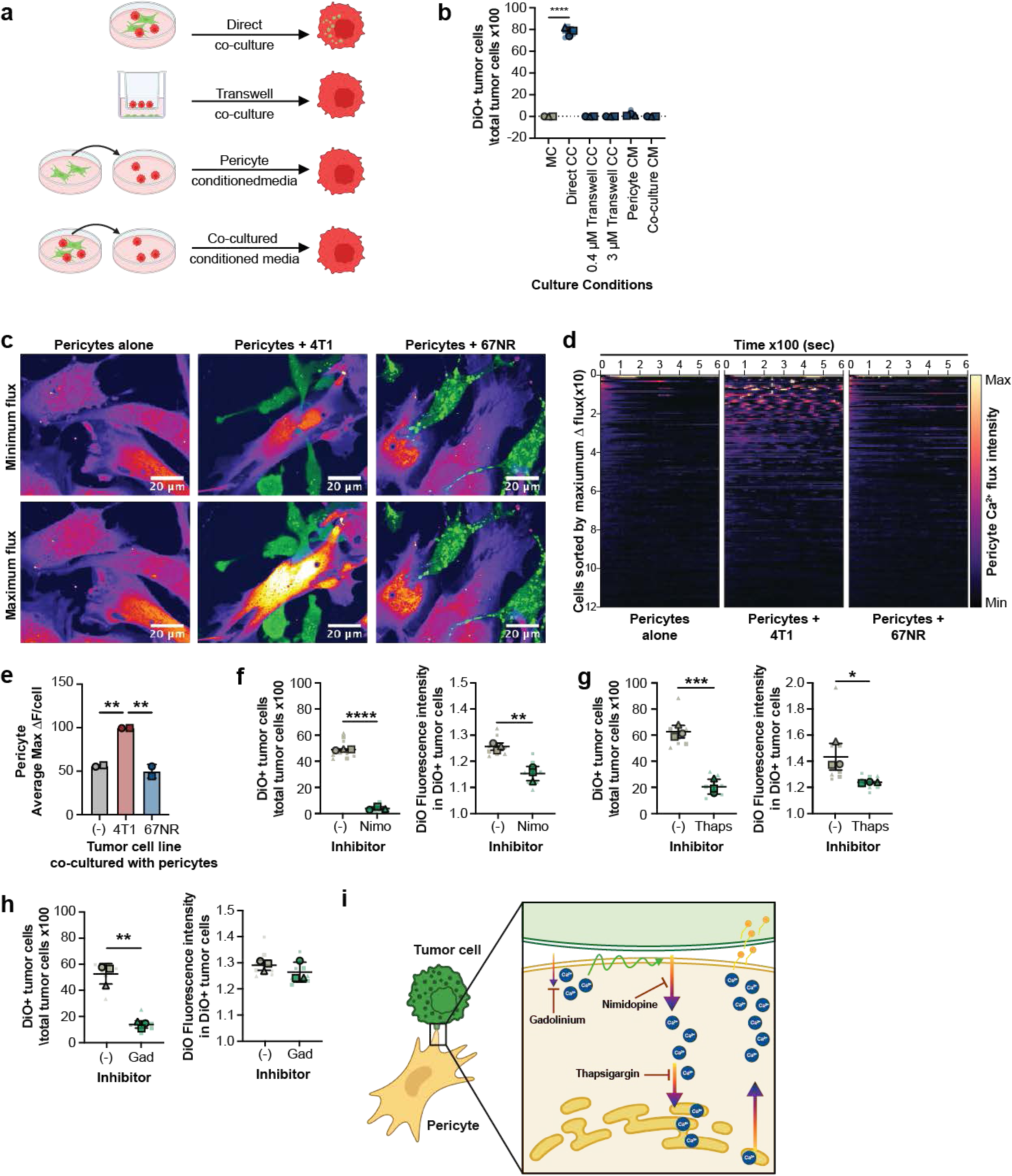
Pericyte Ca2+ channel activation promotes phospholipid transfer. **(a)** Experimental workflow: lipid transfer to tumor cells was assessed in direct co-culture, transwell assays, incubation with conditioned medium from monoculture pericytes, and conditioned medium from direct co-culture. **(b)** The percentage of DiO+ tumor cells in each experimental condition is quantified (one-way ANOVA with Dunnett’s multiple comparisons test; ****p<0.0001). **(c)** Representative images of calcium flux in pericytes during monoculture or co-culture with 4T1 and 67NR cells (green). **(d)** Heat map of raw ΔF/F calcium traces for individual pericytes from monoculture, 4T1 co-culture and 67NR co-culture conditions. **(e)** Quantification of the maximum change in calcium fluorescence intensity per pericyte in each condition (One-way ANOVA with Tukey’s multiple comparisons test, **p<0.01). Quantification of DiO+ tumor cells as a percentage of the total tumor cell population and DiO fluorescence intensity in the DiO+ subpopulation in the presence of **(f)** nimodipine, **(g)** thapsigargin, and **(h)** gadolinium, compared to vehicle control (Unpaired t test, *p<0.05, **p<0.01, ***p<0.001, ****p<0.0001). **(i)** Proposed transfer mechanism: direct contact between pericytes and tumor cells activates mechanosensitive and voltage-gated calcium channels in pericytes, leading to an increase in intracellular calcium. This calcium influx triggers release of pericyte-derived phospholipids, which are subsequently taken up by the adjacent tumor cells.

Pericytes exert homeostatic control over microvascular capillary beds through direct communication with an interconnected network of endothelial, neuronal, and perivascular cells *in vivo*^46–49^. Direct contact-mediated intercellular communication is initiated through an array of related mechanisms, including NOS2, COX2, and prostaglandin pathways, and calcium signaling^47^. We hypothesized that metastatic tumor cells may co-opt pericyte-specific mechanisms of communication during extravasation and colonization. To test this, we utilized inhibitors of these mechanisms during transient pericyte-tumor cell co-culture and examined the percent of tumor cells that received phospholipids, and the degree of phospholipids transfer **(Supplemental Fig 3B)**. Inhibitors of the NOS2, COX2 and prostaglandin pathways had no impact on pericyte-derived phospholipid transfer to tumor cells, nor pericyte or 4T1 tumor cell number **(Supplemental Fig 3C-D)**. By contrast, calcium depletion reduced the amount of pericyte-derived phospholipid that was transferred to 4T1 tumor cells, highlighting a potential role for calcium signaling in this process **(Supplemental Fig 3E)**.

Calcium signaling is critical for perivascular cell responses to mechanical forces such as changes in blood pressure through electrical coupling (syncytia) with neighboring cells, including endothelial cells, vascular smooth muscle cells and other pericytes^48,50–52^. We observed higher numbers of pericytes exhibiting calcium flux, and with larger magnitudes, during transient co-culture with 4T1 tumor cells, suggesting that direct contact with tumor cells promotes calcium channel activation in pericytes **(Supplemental Fig 3F)**. To investigate whether the differential ability of metastatic tumor cells to acquire phospholipids is associated with their enhanced capacity to activate pericyte calcium channels, we performed calcium imaging of pericytes co-cultured with 4T1 and 67NR tumor cells. While 4T1 cells effectively stimulated calcium activity in pericytes, non-metastatic 67NR cells were significantly less effective **(Fig 3C-E, Supplemental Movie 1-9)**. This suggests that metastatic cells have a distinct ability to trigger calcium signaling in pericytes, facilitating the release of phospholipids. Additionally, the ability of 4T1 cells to trigger calcium activity in pericytes appears to be contact dependent as secreted factors from 4T1 cells did not elicit the same response **(Supplemental Fig 3G)**.

Consistent with this hypothesis, inhibitors of electrochemical signal propagation via activation of voltage-gated calcium channels and sarcoplasmic/endoplasmic reticulum calcium pumps, including nimodipine and thapsigargan, decreased the amount and efficiency of pericyte-to-tumor cell phospholipid transfer **(Fig 3F-G)**. L-type voltage-gated channels, such as those that are inhibited by nimodipine, are activated by membrane depolarization events which increase contractile tone in pericytes and smooth muscle cells in response to mechanical signals such as changes in intraluminal pressure^53,54^. Membrane depolarization can occur in response to both local secretory cues and mechanical perturbations^55,56^. Together these data indicate that 4T1 tumor cells initiate phospholipid transfer at least in part by co-opting pericyte calcium-dependent electrochemical propagation machinery.

To investigate the role of direct pericyte-tumor contact dependent mechanosensation as a mechanism for phospholipid transfer, we utilized gadolinium to uncouple membrane deformation from stretch-activated channel activation. Inhibition of mechanosensitive calcium channel activation using gadolinium decreased the percent of tumor cells that received pericyte-derived phospholipids during transient pericyte contact compared to vehicle control-treated cells **(Fig 3H)**. The degree of phospholipids transfer was unchanged in tumor cells that were still able to accept lipid transfer under gadolinium treatment, implying that these specialized force-sensing channels are critical for transfer initiation rather than transfer efficiency **(Fig 3H)**. Together, these observations propose that direct contact induces membrane depolarization events, including through membrane deformation, that in turn promote the transfer of pericyte-derived phospholipids to tumor cells **(Fig 3I)**.

### Pericyte mechanosensitive channel activation is sufficient to promote tumor cell quiescence

Our data demonstrate that transient pericyte contact selectively transfers phospholipids to metastatic tumor cells which promotes tumor quiescence **(Fig 2-3)**. We also observed that calcium channel activation is required for pericyte-tumor cell phospholipid transfer, including through membrane deformation **(Fig 3H)**. Multiple calcium channels exhibit mechanosensitive properties and are activated through mechanical forces such as alterations in blood flow/pressure^57^. We hypothesized that calcium flux in pericytes is initiated by tumor cells through mechanosensitive calcium channels. Gene expression analysis of primary lung pericytes revealed expression of several mechanosensitive calcium channels, of which *Piezo1* was the most highly expressed **(Supplemental Fig 4, Supplemental Table 3)**. We observed greater calcium flux when pericyte cultures were treated with the highly specific stretch-activated PIEZO1 agonist, Yoda1^58^, demonstrating that primary pericyte cultures retain PIEZO1 channel activity *in vitro* **(Fig 4A-C)**.

**Figure 4.**
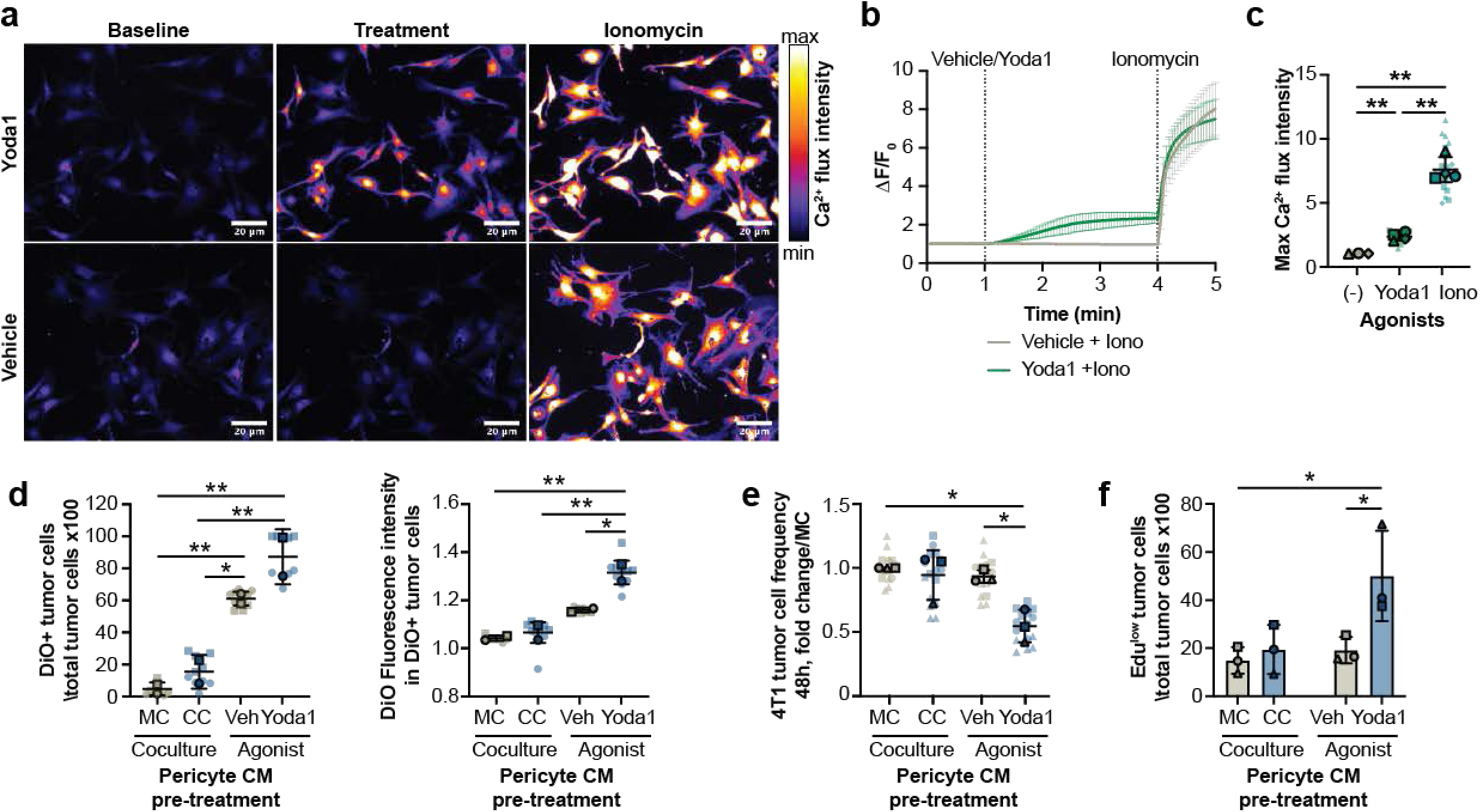
Pericyte Piezo1 Ca2+ channel activation promotes phospholipid transfer and tumor dormancy. **(a)** Representative images of calcium activity in pericytes treated with vehicle or Yoda1, followed by ionomycin. **(b)** Calcium flux trace over time in pericytes treated with vehicle or Yoda1. **(c)** Quantification of maximum calcium flux intensity (One-way ANOVA with Dunnett’s multiple comparisons test, **p<0.01). **(d)** Quantification of DiO+ 4T1 cells as a percentage of the total tumor cell population and DiO fluorescence intensity in the DiO+ subpopulation 2 h following treatment with conditioned medium from 20 min monoculture and co-culture, or pericytes treated with Yoda1 or vehicle for 20 min (One-way ANOVA with Tukey’s multiple comparisons test, *p<0.05, **p<0.01). Tumor cell frequency (One-way ANOVA with Tukey’s multiple comparisons test, *p<0.05, **p<0.01) **(e)** and percentage of EdU^low^ tumor cells **(f)** 48 h following transient monoculture or co-culture, or 20 min incubation with conditioned medium from vehicle and Yoda1 treated pericytes (One-way ANOVA with Bonferroni’s multiple comparisons test, *p<0.05).

The PIEZO1-specific agonist Yoda1 was utilized to activate calcium flux in the absence of direct pericyte-tumor contact to assess the role of pericyte mechanosensitive channel activation in promoting tumor quiescence. We observed higher numbers of 4T1 tumor cells containing pericyte-derived phospholipids when treated with the conditioned medium of Yoda1-treated pericytes **(Fig 4D)**. PIEZO1 activation in pericytes thus appears to be sufficient to promote phospholipid transfer to tumor cells. Consistent with our hypothesis that mechanosensitive pericyte calcium flux promotes tumor cell quiescence, transient exposure to the conditioned medium of Yoda1-treated pericytes resulted in fewer 4T1 tumor cells after 48 hours of culture and concomitant elevated number of tumor cells with low EdU incorporation **(Fig 4E-F)**. Together, these data demonstrate that pericyte mechanosensitive channel activation is sufficient to promote pericyte-to-tumor cell phospholipid transfer resulting in tumor cell quiescence.

### Transient pericyte contact induced tumor dormancy gene signature is associated with metastatic latency in patients

Our data indicate that pericyte-derived phospholipids may promote DTC quiescence. To examine an association between DTC dormancy and phospholipid biology in patients, we compared the gene expression of dormant and proliferative DTCs recovered from bone marrow aspirates of patients with prostate cancer^59^. Similarly to breast cancer, prostate cancer patients also experience a high rate of late metastatic recurrence, indicating the presence of a latent subpopulation of tumor cells that survive primary disease treatment^2,3^. As anticipated, pathway and functional redundancy analysis revealed the enrichment of gene sets associated with cell cycle regulation in dormant DTCs when compared to proliferative DTCs **(Fig 5A, Supplemental Table 4)**. In addition, we observed an enrichment in gene sets associated with lipid and phospholipid metabolism, in keeping with the hypothesis that phospholipid biology is associated with tumor dormancy in patients with metastatic spread **(Fig 5A, Supplemental Table 4)**.

**Figure 5.**
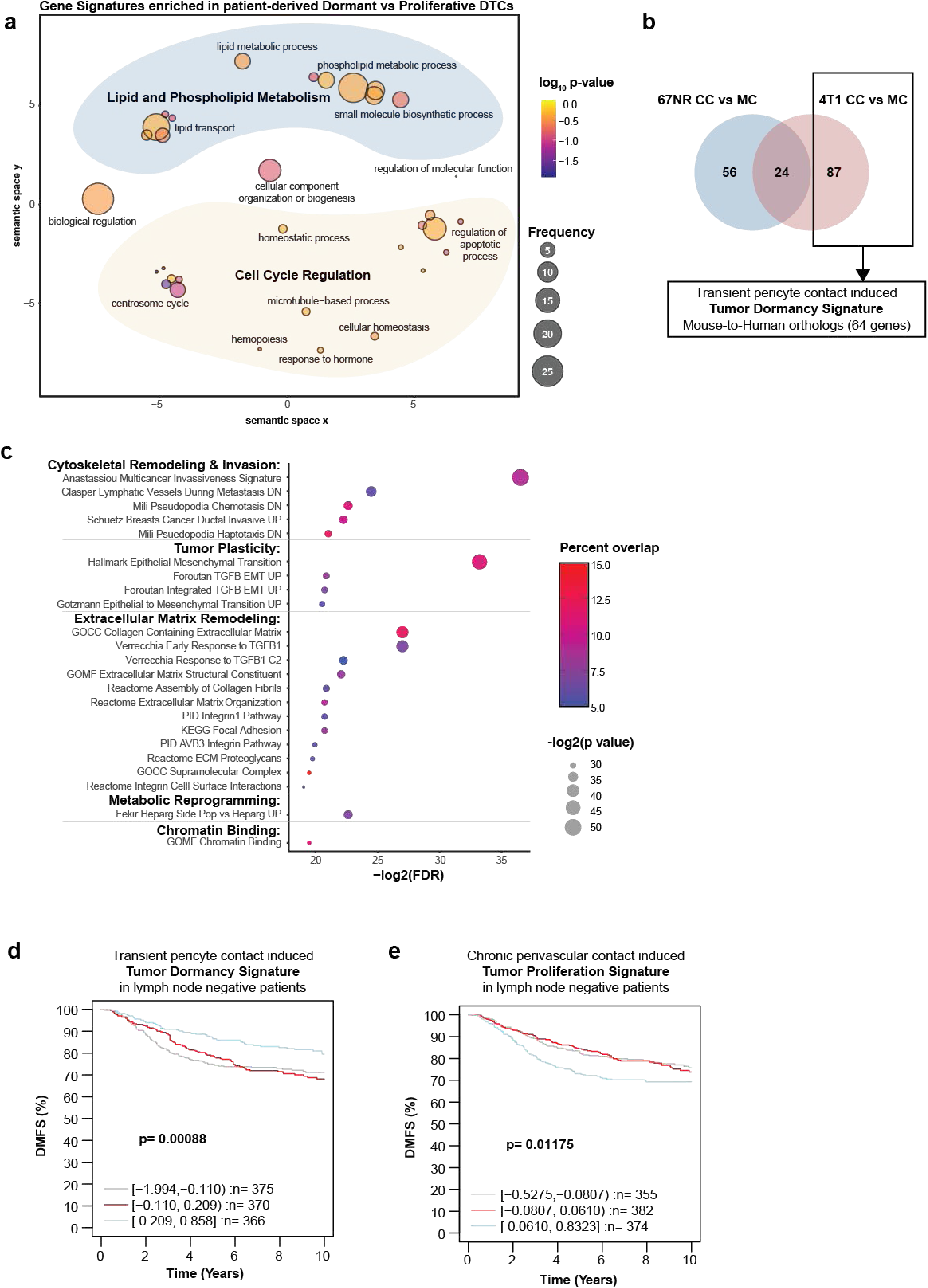
Transient pericyte contact induced transcriptional changes are associated with metastatic latency. **(a)** Enriched gene signatures in dormant vs proliferative DTC recovered from bone marrow of patients with prostate cancer. **(b)** The transient pericyte contact induced dormancy signature was derived from the 87 genes identified as unique to the 4T1 response to 20 min co-culture. **(c)** Percentage overlap between pericyte-induced tumor dormancy signature and published signatures available from the molecular signatures database. **(d)** Distant metastasis-free survival (DMFS) outcome in breast cancer patients with tumor gene expression enrichment of the transient pericyte contact induced tumor dormancy signature. **(e)** DMFS outcome in breast cancer patients with tumor gene expression enrichment of a chronic perivascular cell contact induced tumor proliferation signature.

Our observations propose that transient pericyte-tumor cell contact induces a quiescent phenotype *in vitro* and *in vivo* **(Fig 1)**. To investigate the hypothesis that pericyte-induced tumor dormancy may be generalizable to metastatic relapse in the clinic, we developed a transient pericyte contact induced tumor dormancy gene signature and compared this signature to metastatic outcome in patients. We first compiled genes that were differentially expressed in 4T1 cells from dormancy-promoting transient pericyte co-culture compared to those from monoculture with a p-value cut off <0.05, which yielded a total of 111 genes **(Fig 5B, Supplemental Table 5)**. We next compared these genes to those that were differentially expressed in 67NR transient co-cultured cells compared to those from monoculture, yielding a total of 80 genes **(Fig 5B, Supplemental Table 6)**. 24 genes were overlapping between 4T1 and 67NR in co-culture compared to their corresponding monoculture controls. 67NR cells did not exhibit quiescence after transient pericyte co-culture (**Fig 1D**). We therefore reasoned that genes that were enhanced in 4T1:pericyte co-culture but not in 67NR:pericyte co-culture would best represent the pericyte-induced tumor dormancy state. 87 genes were identified as differentially expressed in 4T1 cells, but not 67NR cells, after transient pericyte co-culture **(Fig 5B, Supplemental Table 7)**. The translation of our mouse-derived tumor dormancy signature to human orthologs yielded a total of 64 genes for downstream analysis **(Fig 5B, Supplemental Table 8)**.

To investigate the relationship between our pericyte-induced tumor dormancy signature and published signatures related to tumor progression, we computed the percent overlap with gene sets available on the Molecular Signatures Database^60^. Only 5-15% overlap was computed between the tumor dormancy signature and published gene sets, indicating that the pericyte-induced tumor dormancy signature may represent a previously undescribed mechanism for tumor dormancy compared to previous studies **(Fig 5C, Supplemental Table 9)**. Low overlapping pathways in this analysis included cellular processes related to cytoskeletal remodeling, tumor plasticity, extracellular matrix remodeling, metabolic reprogramming and chromatin binding, suggesting an overlapping relationship between known mechanisms of metastatic relapse and pericyte-induced tumor dormancy (**Fig 5C, Supplemental Table 9**).

To probe the relationship between pericyte-induced tumor dormancy and metastatic outcome, we computed the enrichment of the transient pericyte contact induced tumor dormancy signature to the gene expression in tumors from breast cancer patients who were profiled when their disease was negative for lymph node metastases^61^. Distant metastasis-free survival (DMFS) was significantly greater in patients whose tumors were highly enriched for the pericyte-induced tumor dormancy signature, revealing a potential relationship between tumor gene expression changes due to transient pericyte contact and metastatic latency **(Fig 5D)**. Conversely, DMFS was significantly reduced in patients with high tumor gene expression enrichment of a chronic perivascular cell contact induced tumor proliferation signature **(Fig 5E, Supplemental Table 10)**, generated from D2A1 tumor cells after chronic co-culture with perivascular smooth muscle cells. Together these results suggest that pericyte-tumor crosstalk may activate tumor cellular programs that are critical for tumor cell fate and metastatic outcome.

## Discussion

Metastatic recurrence is the leading cause of cancer related deaths^1^. It is increasingly recognized that tumor cell dissemination is initiated early in primary tumor development^4–7^. While most of these disseminating tumor cells (DTCs) are cleared by the immune system, some survive and persist in a dormant state, below clinical detection^10,62^. These dormant cells act as reservoirs for metastatic relapse, which can occur months to decades after successful treatment of the primary tumor^3,63^. The reliance of DTC on local stromal cells to aid survival and colonization has been extensively reported, including in models of breast cancer^13,14,21^. Here, we demonstrate that pericytes in the lung microenvironment facilitate the induction of DTC dormancy, a previously unappreciated mechanism in the early metastatic niche.

Previous work has shown that pericytes remodel the extracellular matrix (ECM) in response to tumor-derived secreted factors, initiating the formation of a pre-metastatic niche by producing fibronectin-rich ECM that promotes metastatic outgrowth^11^. However, direct interactions between tumor cells and pericytes, particularly outside the CNS, remains understudied. Given that pericytes are located along blood vessels^46^ and are among the first stromal cells encountered by tumor cells during extravasation, we sought to understand the consequences of pericyte-tumor cell contact on metastasis. Our findings reveal a novel pericyte-dependent mechanism for DTC dormancy that is mediated by the transfer of pericyte-derived lyso-PLs. This transfer is initiated by transient tumor cell activation of pericyte calcium channels, resulting in the release of lyso-PLs that promote tumor cell quiescence. This previously unappreciated role of pericytes in promoting DTC latency highlights new therapeutic opportunities to prevent metastatic relapse.

In agreement with our previous studies, our experiments confirm that chronic exposure to pericytes promotes tumor proliferation, likely through ECM remodeling or altered secretion of growth factors^11^. Surprisingly, our results also indicate that transient contact with pericytes, such as that which would occur during extravasation, instead results in dormancy induction. We hypothesize that pericytes initially promote dormancy to protect DTCs from microenvironmental stress but may later facilitate reawakening by remodeling the niche under specific signals. These findings highlight the multifaceted role of pericytes in metastasis and suggest that their influence on metastatic tumor progression depends on the timing and duration of tumor-pericyte interactions. While this work focused on the lung, the mechanism appears relevant in the brain and may represent a cancer-type-agnostic pathway which could have far reaching clinical benefit.

Pericytes have a vital function in regulating blood flow by responding to calcium signaling and electrical coupling with neighboring endothelial and other mural cells. Calcium signaling not only regulates vascular tone by contractile machinery^64^ but also mediates the release of vasoactive molecules^65^, including lysophosphatidic acid (LPA), essential for vascular homeostasis^65,66^. Our study identified the functional expression of PIEZO1, a mechanosensitive cation channel^67^, on primary lung pericytes. PIEZO1 activation facilitated phospholipid transfer to DTCs, promoting their dormancy. While others have reported *Piezo1* expression on brain capillary pericytes^68^, to our knowledge its presence on pericytes outside the CNS has not been documented. PIEZO1 has a known role in mediating the endothelial responses to fluid shear stress and blood flow regulation^69,70^, its expression in pericytes may contribute a second level of control over blood flow that has not yet been fully explored in addition to a role in phospholipid-induced tumor dormancy. Our findings reveal that pericytes leverage physiologically relevant calcium signaling to facilitate the transfer of lyso-PLs to tumor cells, promoting their persistence in the metastatic niche.

Metabolomics analysis revealed enrichment of specific lyso-PLs in tumor cells following transient pericyte contact, three of which were annotated. Lyso-PLs play critical roles in regulating cell survival and proliferation across various pathologies, including atherosclerosis, inflammation, and cancer^66,71,72^, where their effects appear to be cell-type and context-specific. For example, lyso-PC inhibits endothelial cell migration and proliferation^73^ but promotes the proliferation of vascular smooth muscle cells^36^ and contributes to pericyte loss in the CNS^74^. In cancer, lyso-PLs also exhibit context-dependent effects. While lyso-PC has exhibited anti-tumor activity^34^ and reduced the growth of macrometastatic lesions in a variety of tumor cells^75^, these studies did not assess the presence or absence of micrometastatic lesions and single dormant DTC. Lyso-PE has been observed to enhance the proliferation and migration of breast cancer cells^71^ and drive migration and invasion in ovarian cancer cells^72^. These contrasting effects may depend on both the type of lyso-PLs and the duration of exposure. Future work will explore the precise mechanisms through which pericyte-specific lyso-PLs influence the cell cycle and induce tumor cell dormancy, which might include direct signaling by lipids as secondary messengers^66,76^, metabolic^24,25,27^, or epigenetic reprogramming^13^.

While the transfer of materials/signaling molecules between stromal and tumor cells has been observed before^42–45,77^, our findings identify pericytes as a previously unidentified source of these materials, through perivascular cell specific mechanisms, and with hitherto unreported outcome. Interestingly, our experiments demonstrated that in a diverse panel of stromal cell populations, pericytes were the most efficient in transferring lipids to tumor cells, followed by vascular smooth muscle cells, both of which were more effective than fibroblasts and macrophages; this indicates that shared perivascular cell functional machinery likely underpins this process. Furthermore, we also identified greater persistence of DTC in the brain after pericyte co-culture, but not in the liver or bone. While a similar mechanism of pericyte driven dormancy appears to aid tumor cell survival in the brain microenvironment, future studies are needed to determine whether differences in liver and bone are reflective of their unique microenvironment or that of our tumor cell delivery model.

Given that chemotherapy targets actively proliferating cells, limiting or even eliminating quiescent DTCs presents a major therapeutic challenge. In this study, we identified calcium channel inhibitors that block lyso-PL transfer between pericytes and tumor cells. These inhibitors are promising, as many are already used clinically for other conditions^78–81^. Notably, drugs in this class are being explored for triple negative breast cancer treatment; for example, Diltiazem, an approved antihypertensive, has been shown to affect motility, EMT phenotype, colony formation, and metastasis in 4T1 cells in mice^78^. Nifedipine has been demonstrated to reduce colorectal cancer proliferation and metastasis by reducing calcium activated NFAT2 transcriptional activation^79^.

Additional therapeutic approaches may target receptor-mediated uptake of lipid signaling molecules, such as the S1P receptor antagonists, Fingolimod^80^ and Siponimod^81^, which are FDA approved for multiple sclerosis. Lyso-PL uptake occurs via multiple mechanisms, including passive diffusion, receptor mediated uptake e.g., LPARs and S1PRs, endocytosis or lipid transport proteins^82–84^. Future work is needed to uncover the lyso-PL uptake mechanism to determine which inhibitors might be most effective in limiting lyso-PL transfer.

This study highlights the importance of tumor-pericyte interactions in directing DTC dormancy. We propose that tumor cells initiate calcium-dependent lyso-PL transfer from pericytes, inducing a dormant state to protect against environmental stressors at the metastatic site until conditions become favorable for regrowth. Future research will focus on understanding how lyso-PLs influence tumor cell cycling and uncovering downstream pathways that regulate dormancy. Targeting these pathways could uncover new therapeutic opportunities to prevent metastatic relapse.

---

**Supplemental Figure 1. (a)** Percentage of tumor cells that interacted with pericytes during extravital imaging of the lung. **(b)** Experimental workflow: Tumor cells were co-cultured with zsGreen+ pericytes for 20 min and then retrieved from the media suspension, collected tumor cells were seeded on a new plate and imaged for contamination of zsGreen+ pericytes. **(c)** Total number and percentage of zsGreen+ pericytes in chronic co-culture vs transient monoculture and co-culture. **(d)** E0771 tumor cell proliferation 48 h after isolation from transient co-culture with pericytes; compared to tumor cell monoculture controls (Unpaired t test, *p<0.05). **(e)** Graph showing intensity of p27k mVenus expression in quiescent cells vs cycling cells over time. **(f)** Percentage of mice with viable 4T1 cells retrieved from the liver, bone and brain, 10 days following intracardiac injection of 4T1 from monoculture or co-culture. **(g)** Percentage of mice with viable 4T1 cells retrieved from lungs at 3 and 10 days following intracardiac injection of tumor cells from monoculture or co-culture. **(h)** Proportion of mCherry+ cells in the lung 3 days after intracardiac injection of tumor cells from monoculture or co-culture. **(i)** Proportion of EdU+ mCherry+ tumor cells of total mCherry positive population in the lungs 3 days after intracardiac injection.

**Supplemental Figure 2. (a)** Metabolic pathways enriched in 4T1 and 67NR cells from transient pericyte co-culture compared to monoculture. **(b)** Volcano plots of differences in metabolites identified in 4T1 co-culture versus 67NR co-culture, and metabolites enriched in 67NR cells exposed to transient pericyte precondition or monoculture control. **(c)** DiO fluorescence intensity in the DiO+ subpopulation (Unpaired t test, ***p<0.001). **(d)** Experimental workflow: Pericytes were grown in the presence of a methionine analogue (HPG) for 1 week prior to co-culture to allow incorporation into pericyte synthesized proteins. mCherry+ 4T1 tumor cells were maintained in methionine (Me) containing medium and then co-cultured with mCherry+ 4T1 cells for 2 h to allow the transfer of pericyte derived HPG labelled proteins and fixed. A click chemistry reaction was performed to identify the transfer of pericyte derived proteins in tumor cells. **(e)** Percentage of HPG positive tumor cells following monoculture or co-culture with pericytes. HPG was added to 4T1 cells for 30 min prior to fixing as a positive control (One-way ANOVA and Sidak’s multiple comparisons test, ****p<0.0001). **(f)** Quantification of cell tracker (CT) green+ tumor cells as a percentage of the total tumor cell population and CTgreen fluorescence intensity in the CTgreen+ subpopulation. **(g)** Quantification of mT/mG+ tumor cells as a percentage of the total tumor cell population and mT/mG fluorescence intensity in the mT/mG+ subpopulation (One-way ANOVA with Sidak’s multiple comparisons test, *p<0.05). **(h)** Representative images of cholesterol transfer from pericytes pre-loaded with bodipy-cholesterol and then co-cultured with DiD labelled 4T1 and 67NR cells for 24 h.

**Supplemental Figure 3. (a)** DiO fluorescence intensity in tumor cells across experimental condition is quantified (one-way ANOVA with Dunnett’s multiple comparisons test; ****p<0.0001). **(b)** Identification of inhibitors to target physiologically relevant pericyte communication pathways. Quantification of DiO+ tumor cells as a percentage of the total tumor cell population, DiO fluorescence intensity in the DiO+ subpopulation, and cell count of pericytes and tumor cells in the presence of **(c)** aminoguanidine or **(d)** indomethacin, compared to vehicle control. **(e)** Quantification of DiO+ tumor cells as a percentage of the total tumor cell population and DiO fluorescence intensity in the DiO+ subpopulation following co-culture in low calcium hanks solution compared to normal calcium containing hanks solution (Unpaired t test, **p<0.01). **(f)** Representative images of calcium flux in pericytes during monoculture and 4T1 (green) co-culture. Heat map of raw ΔF/F calcium traces for individual pericytes from monoculture and 4T1 co-culture. Quantification of the maximum change in calcium fluorescence intensity per pericyte in each condition (Mann Whitney test, ***p<0.001). Percent of pericytes exhibiting calcium flux above a set threshold defined as the average of monoculture calcium flux plus the standard error (Fishers exact test, p= 0.0087). **(g)** Average maximum change in calcium fluorescence intensity per pericyte in monoculture, co-culture and 4T1 TCM conditions (One-way ANOVA with Tukey’s multiple comparisons test, *p<0.05).

**Supplemental Figure 4.** Heat map of transcript counts of various ion channels from three batches of primary lung pericytes.

## Materials and Methods

### Animals

For injection of BALB/c derived tumor cell lines, BALB/cJ mice were purchased from The Jackson Laboratory and injected at 10-12 weeks old. NG2-ERT-creT2 ROSA-STOP-flox-zsGreen mice, also referred to as NG2 lineage tracing mice were used to track pericytes *in vivo* and were maintained under pathogen free conditions within the NIH animal facility. Genotyping was performed via PCR using validated protocols. For euthanasia, all animals were anesthetized by isoflurane followed by cervical dislocation. All animal experiments were conducted in accordance with the guidelines and regulations outlined in the Guide for the Care and Use of Laboratory Animals and were approved by the NCI-Bethesda Animal Care and Use Committee. All procedures were in accordance with those outlined in the ARRIVE guidelines.

### Extravital Imaging

Intraperitoneal injections of tamoxifen (T5648, Millipore Sigma) were administered to induce CreERT-mediated recombination. Tamoxifen was dissolved in corn oil (C8267, Millipore Sigma) to a final concentration of 10 mg/mL and 100 µL was injected once a day for 10 d total (5 d on, 2 d off, 5 d on) followed by a one-week wash-out period. NG2-pericyte zsGreen lineage tracing mice were euthanized by cervical dislocation and immediately intracardiac injected with 1×10^5^ mCherry expressing 4T1 cells and the vascular tracer 5-(and-6)-tetramethylrhodamine biocytin (Biocytin TMR; T12921, Thermo Fisher Scientific) used at 10 mM in PBS (T12921, Thermo Fisher Scientific). Collected lungs were imaged immediately using a Nikon SoRa Spinning Disk confocal microscope. Images were processed and analyzed using Imaris software (v9.9.0), focusing on the association between tumor cells and pericytes within a 2D distance categorized into bins of less than or greater than 20 µm from the nearest pericyte soma, at greater than or less than 20 min from the beginning of imaging.

### Metastasis Assay

Mice were anesthetized with isoflurane (NDC 10019-360-40, Baxter Healthcare Corporation) and positioned in lateral recumbency. A 28-gauge needle (329461, BD) was inserted into the chest perpendicular to the bench top. The syringe was gently aspirated to confirm correct placement, indicated by a flash of blood. The 4T1 cell suspension collected from monoculture or co-culture, containing 1×10^5^ cells, was injected in 100 µL of Hanks’ balanced salt solution (HBSS; 14175103, Thermo Fisher Scientific).

After 10 days, mice were anesthetized with isoflurane and euthanized by cervical dislocation. Tissues were cardiac perfused with PBS to remove blood, dissected and placed in ice-cold PBS. The left lung lobe was collected for confocal imaging and live lung surface imaging was performed to identify mCherry expressing 4T1 lesions. The remaining lobes were processed for dissociation. Lung tissue was cleaned to remove connective tissue, minced and digested in 1 mg/mL collagenase I (17100-017, Thermo Fisher Scientific), 10 µg/mL Dispase II (4942078001, Millipore Sigma), 20 µg/mL DNase I (D4263, Millipore Sigma), containing HBSS and incubated at 37°C for 20 min in a shaking heat block.

Livers, with gallbladders removed, were minced and digested at 37°C for 50 min. Every 15 min, tissue was further dissociated with a pipette tip, the suspension was collected, and an equal volume of fresh digestion medium was added. Collected suspensions were kept on ice.

Femurs were collected and gently crushed with a mortar and pestle in 1 mL of HBSS containing 2% FBS and 1 mM EDTA (15575020, Thermo Fisher Scientific). This was repeated 3 times to ensure maximum marrow cell collection.

Brains, excluding brain stem, cerebellum and olfactory bulb, were rolled on blotting paper to remove meninges, minced and digested in collagenase A (11088793001, Millipore Sigma) and DNase1 at 37°C for 40 min with agitation. 20% BSA (9048468, Gold Biotechnology)-DMEM was used to remove myelin.

All digested tissues were filtered and plated on 10 cm dishes in selection medium containing RPMI (11-875-119, Thermo Fisher Scientific), 10% Fetal Bovine Serum (FBS; 900108500, GeminiBio), 1% penicillin/streptomycin (15070063, Thermo Fisher Scientific) and 60 µM 6-Thioguanine (B21280.06, Thermo Fisher Scientific). Selection medium was refreshed every 2–3 days. Outgrowing colonies were imaged to confirm mCherry expression using the EVOS FL Auto 2 Cell Imaging System (AMAFD2000, Thermo Fisher Scientific).

### Primary cell isolation and culture

Primary lung pericytes were isolated as previously described^85^ from 12-week-old female C57BL/6 mice or mT/mG reporter mice^86^ and maintained at 37°C, 5% CO_2_ and 10% O_2_ (physiological oxygen for lung pericytes^85^) on collagen coated plates. Pericytes were grown in medium containing a 1:1 mix of low glucose Dulbecco’s Modified Eagle’s medium (DMEM) (D6046, Millipore Sigma) and Ham’s F12 nutrient mix (11765047, Thermo Fisher Scientific), supplemented with 1% MEM non-essential amino acids (11140050, Thermo Fisher Scientific), 2mM sodium pyruvate (11360070, Thermo fisher Scientific), 20mM Hepes (15630080, Thermo Fisher Scientific), 1X endothelial cell growth factors (ECGF; 390599, Bio-techne), 1% penicillin/streptomycin and 20% FBS (900-208-500, GeminiBio). Before experiments, pericytes were placed in growth arrest (GA) medium containing reduced FBS (2%) and no ECGF for 24 h. All experiments were performed with passage 1 pericytes.

Primary dermal fibroblasts, provided by Christophe Cataisson and isolated as previously described^87^, were maintained in DMEM (10313021, Thermo Fisher Scientific), supplemented with 10% FBS, 1% glutamine (25030081, Thermo Fisher Scientific) and 1% penicillin/streptomycin.

Primary vascular smooth muscle cells (VSMCs) were isolated from mouse aortas. Briefly, aortas were dissected, cleaned, and cut into small fragments, then digested in DMEM with 1.42 mg/mL collagenase II (17101015, Thermo Fisher Scientific) for 6 hours at 37°C. The cell suspension was centrifuged, resuspended in DMEM supplemented with 10% FBS, 1% glutamine and 1% penicillin/streptomycin, and plated in 24-well plates for 5 days to allow VSMC outgrowth.

Primary macrophages were generated from murine bone marrow as previously described^88^. Bone marrow was collected from the femurs of mice by crushing using a mortar and pestle. RBC were removed using ACK lysis buffer (10-548, Lonza), cells were filtered, counted and plated in pericyte basal medium supplemented with 15% L929 CM and 1% penicillin/streptomycin for 5 days, followed by pericyte basal medium, 1:50 ECGF, 15% FBS, 1:1000 macrophage-colony stimulating factor (300-332P, GeminiBio) for 5 days. L929 CM was generated from 80% confluent cells in DMEM, 10% FBS, 1% penicillin/streptomycin and 1% glutamine, over 10 days.

### Immortalized cell lines and culture

4T1 cells were provided by Rosie Kaplan (National Cancer Institute, Bethesda, USA) and grown in RPMI with 10% FBS and 1% penicillin/streptomycin. 4T07, 67NR, D2.0R and D2A1 cells were provided by Kent Hunter (National Cancer Institute, Bethesda, USA) and grown in DMEM containing 10% FBS, 1% glutamine and 1% penicillin/streptomycin. The immortalized human non-tumor mammary epithelial cell line MCF10A (M1) and the metastatic counterpart MCF10 Ca1a.c11 (M4), were provided by Lalage Wakefield (NCI, Bethesda, USA). M1 cells were grown in DMEM/F12 containing 5% horse serum (H1138, Millipore Sigma), 10 μg/mL Insulin (I6634, Millipore Sigma), 20 ng/mL epidermal growth factor (300-110P, Gemini Bio), 500 ng/mL Hydrocortisone (H0888, Millipore Sigma), 100 ng/mL Cholera Toxin (C8052, Millipore Sigma), 1% penicillin/streptomycin. M4 cells were grown in DMEM/F12 containing 5% horse serum, and 1% penicillin/streptomycin. Immortalized human lung pericytes^89^ were provided by Hongkuan Fan (University of South Carolina, Charleston, USA) and cultured on 0.2% gelatin (G1393, Millipore Sigma) coated plates in commercial pericyte medium (1201, ScienCell Research Laboratories). All immortalized cell lines were maintained at 37°C humidity-controlled incubator with 5% CO_2_.

### Lentiviral production and transduction

To measure quiescence, we used a previously published mVenus-p27K reporter^23^, consisting of mVenus fused to an inactive p27-CDK binding domain expressed by the ubiquitous promoter PGK. The mVenus-p27K plasmid was provided by Judith Agudo (Dana-Farber Cancer Institute, Harvard Medical School Boston USA). The plasmid was eluted from filter paper into TE buffer and transformed into Shot TOP10 chemically competent bacteria, as per TOP10 instructions. Competent bacteria were provided by Steve Cappell (National Cancer Institute, Bethesda, USA). Transformed bacteria were spread on Carbenicillin plates (11006443, IPM Scientific) and allowed to incubate at 37°C overnight, and then selected colonies were expanded in shaking liquid culture overnight at 37°C. Plasmid DNA was isolated via Midiprep performed using the Qiagen Plasmid Plus Midi Kit (12943, Qiagen) as per the manufacturer’s instructions. Plasmid quality and concentration were confirmed via Nanodrop spectrophotometry.

Using the isolated plasmid, third generation lentiviral particles were produced through transfection into 50% confluent HEK293T cells cultured in serum-free Opti-MEM medium (31985070, Thermo Fisher Scientific). Supernatants were collected daily for 72 h, passed through a 0.45 µm filter, concentrated at 3600 rpm for 20 min in 15 mL Amicon Ultra tubes (UFC910024, Millipore Sigma) with a 100,000 kDa MW cutoff, and frozen at -80°C in aliquots. For lentivirus transduction into cells of interest, tumor cells were cultured with lentivirus in medium containing 5µg/mL polybrene (TR-1003, Millipore Sigma) at 37°C for 16-20 h. Medium was replaced, and cells were expanded and subsequently selected for with puromycin (A1113803, Thermo Fisher Scientific) for two weeks. Reporter function was validated through proteasome inhibitor (MG-132; M7449, Millipore Sigma) treatment and imaging.

### Transient Co-culture assays

Pericytes were seeded in 12-well plates at 5×10^4^ cells per well in GA medium overnight. The next day tumor cells are trypsinized, resuspended in GA medium and added to pericyte culture at a 1:1 ratio. After 20 min co-culture, tumor cells are gently isolated from the suspension of the co-culture (pericytes are adhered but tumor cells are not in this time frame). Tumor cell suspension was centrifuged and resuspended in the appropriate tumor cell medium and seeded for downstream analysis.

### Chronic Co-culture assays

Pericytes (5 × 10³ cells) were seeded in GA medium onto black, clear-bottom 96-well plates (3598, Corning) and allowed to adhere overnight. The following day, tumor cells (5 × 10³ cells) were added to the pericytes and co-cultured for 48 h. For experiments involving unlabeled tumor cells, tumor cells were pre-incubated with CellTracker Green (C2925, Thermo Fisher Scientific) for 40 min at 37°C before seeding. At the experiment endpoint, 2 μM Hoechst 33342 nuclear dye (62249, Thermo Fisher Scientific) in PBS was applied to cells. Cell counting was performed using the Lionheart FX automated imager (LFXW-SN, Agilent), where tumor cells were identified as nuclei positive for mCherry (4T1) or CellTracker Green (67NR) using Agilent BioTek Gen5 software (v3.12).

### Live imaging co-culture

For live co-culture imaging, lung pericytes were seeded at a density of 5×10^3^ cells/well in a 96-well black clear bottom plate in GA medium and allowed to adhere overnight. The following day pericytes were labelled with either 10 μM DiO lipophilic dye (60011, Biotium), 5 μM CellTracker Green, or 1 μM Bodipy-Cholesterol (HY-125746, MedChemExpress) in GA medium for 40 min in a 37°C incubator. After labeling, the dye solution was gently removed and cells washed with PBS before replacing with GA medium.

Tumor cells were trypsinized, counted and resuspended in pericyte GA medium. If tumor cells were unlabeled, they were incubated with 10 μM DiD lipophilic dye (D7757, Thermo Fisher Scientific) at 37°C for 40 min before co-culture. The 96-well plate containing the labelled pericytes was transferred to the Lionheart FX imager with humidity, temperature, and O_2_-CO_2_ controls. For imaging, labelled tumor cells were added at a 1:1 ratio and imaged at set time points.

### Soft Agar Colony Forming Assay

Tumor cells were treated with 50 uM Lyso-PC 18:1 (845875P, Avanti Research) or vehicle (ethanol) in RPMI containing 10% FBS and 1% penicillin/streptomycin for 20 min. Tumor cells were collected, centrifuged and seeded at 100 cells/well in a 48-well plate in 200 uL of RPMI medium containing 10% FBS and 1% penicillin/streptomycin and 0.3% agar (A9045, Sigma Aldrich) overlaid onto 300 uL of RPMI medium containing 10% FBS, 1% penicillin/streptomycin and 0.8% agar. Images of colonies were obtained after 7 days using the lionheart FX imager and analyzed in Gen5.

### Immunofluorescence

For immunohistochemistry, at the time of collection the tissues were cardiac perfused with PBS followed by 4% paraformaldehyde (PI28906, Fisher Scientific). Tissues were then harvested and fixed overnight in 4% paraformaldehyde at 4°C. Lungs were then transferred to 30% sucrose/PBS and incubated at 4°C for 3 days. The tissues were embedded in cryomolds (25608-916, Avantor) containing O.C.T. Compound (Optimal Cutting Temperature; 23-730-571, Fisher Scientific) and frozen using methanol in dry ice, and stored at -80°C for cryosectioning. Before staining, sections were air-dried for a minimum of 2 h, washed in PBS, permeabilized in 0.5% Triton-X (X100, Millipore Sigma), blocked in 3% fish gelatin (G7765, Sigma Aldrich)/10% donkey serum (S30-M, Millipore Sigma), incubated with 0.1% sudan black (199662, Millipore Sigma) in 70% ethanol to quench tissue autofluorescence and incubated with primary antibodies overnight at 4°C. The following primary antibodies were used: rabbit anti-Ki67 (ab16667, Abcam), rat anti-mCherry (M11217, Thermo Fisher), and mouse anti-Actin, α-smooth muscle FITC conjugated (F3777, Millipore Sigma). The following day the cells were washed with 0.1% Tween20 (655204, Millipore Sigma) before incubation with donkey secondary antibodies (Jackson ImmunoResearch) for 1 h at room temperature, protected from light. The cells were washed again in 0.1% Tween20 solution followed by incubation with DAPI (D1306, Thermo Fisher Scientific) and finally mounted with ProLong™ Gold Antifade mountant (P36930, Thermo Fisher Scientific). Images were obtained on the Nikon SoRa spinning disk microscope and analyzed in Image J or the Zeiss Axioscan 750 slide scanner and analyzed in Halo. The Halo Quantitative Pathology platform by indica labs was used for segmentation and fluorescence intensity quantification of confocal images by blinded reviewer. The CytoNuclear FL v2.0.12 algorithm was used to segment cells with the following parameters: Nuclear Contrast Threshold, 0.51; Minimum Nuclear Intensity, 0; Nuclear Size, 3571.1; Minimum Nuclear Roundness, 0; Nuclear Segmentation Aggressiveness, 1; Fill Nuclear Holes, True; Maximum Cytoplasm Radius, 5. Single cell fluorescence intensity measures were exported and analyzed using in RStudio. Fluorescence intensity thresholds to identify mCherry+, Ki67+, and Edu+ cells were set using Secondary only, and single-stain controls.

### EdU Incorporation

To label proliferating cells *in vivo*, 200 µg of 5-Ethynyl-2’-deoxyuridine (EdU; CCT-1149, Vector Laboratories) in 100 µL was administered via intraperitoneal injection 16 h prior to euthanasia. For *in vitro* proliferation analysis, EdU was added to culture medium at a final concentration of 10 µM, 16 h prior to cell collection. Cells were fixed with 4% paraformaldehyde for 7 min at room temperature, washed with 3% BSA in TBS (28358, Thermo Fisher Scientific), and permeabilized with 0.5% Triton X-100 for 20 min. The EdU detection was performed using a click chemistry reaction cocktail containing 1x TBS, 4 mM Copper(II) sulfate (Acros, 197730010), 1 µM azide (Click Chemistry Tools, 1275-1), and 100 mM sodium ascorbate (Acros, 352685000) for 30 min at room temperature, followed by a nuclear counterstain.

### Transwell assays

Transwell assays were performed using 24-well 0.4 µm (38024, Costar) and 3.0 µm pore transwell inserts (353096, Falcon). A total of 5×10^4^ pericytes in 500 µl of GA medium were added to the transwell insert which formed the top chamber. 5×10^4^ tumor cells were seeded on a 24-well plate in GA medium which formed the lower chamber. Both cell lines were allowed to adhere overnight to the respective compartment. The following day the pericytes in the top chamber were incubated with DiO for 40 min in GA medium. The pericytes were washed with PBS to remove any unbound dye and GA medium was replaced. The top chamber was then added to the plate with the tumor cells and incubated in a humidified incubator with 5% CO_2_ and 10% O_2_ for 24 h to assess exchange of lipid dye without direct cell contact. The transwell insert was removed and the bottom chamber with tumor cells was imaged for the presence of DiO co-localized with mCherry expressing tumor cells using the lionheart FX imager.

### RNA sequencing

RNA-Sequencing was performed by the CCR-Sequencing Facility on pericytes from GA conditions or tumor cells following isolation from a 20 min co-culture with pericytes or from monoculture as a control. RNA was extracted from pericytes seeded at 1×10^5^ in a 12-well plate or from tumor cells in a cell pellet after isolation from monoculture or co-culture using the RNeasy Mini Kit (74104, Qiagen), quantified and stored at -80°C. Three replicates per experimental condition were collected. RNA-Sequencing was performed on pooled samples to obtain 2×100 bp paired end reads, by the NCI Sequencing facility using the Illumina NextSeq 2000 P2 platform.

Data processing and statistical analysis was performed using Partek flow. Reads were trimmed to remove adapters and low-quality bases using Cutadapt. Minimum read length was set at 25 and minimum Phred quality score of 20. The trimmed reads were aligned with the reference genome (mm10) using STAR. The transcript abundance was estimated using the Partek E/M algorithm. Gene counts were normalized to counts per million and differential expression analysis was performed using DESeq2. For the analysis of co-culture vs monoculture, a p-value cut off of 0.05 was applied due to the small gene sets obtained.

A pericyte-induced tumor dormancy signature was derived from DEG that were unique to 4T1 co-culture condition, but not 67NR co-culture **(Supplementary Table 8)**. Similarly, a perivascular cell-induced tumor proliferation gene signature was generated using D2A1 tumor cells following chronic co-culture with MOVAS cells **(unpublished; Supplementary Table 10).** For each signature the fold change values of the gene were used as the weight vector. Mouse genes were mapped to their human homologs to enable application of the signatures to human datasets. Kaplan-Meier survival analyses were performed using the GOBO web application^61^.

### Analysis of publicly available datasets

The GEO dataset GSE48995 contains microarray expression data from disseminated prostate cancer tumor cells isolated from bone marrow samples. These samples were categorized into 3 subgroups: advanced disease (ADV_1 and ADV_2) or no evidence of disease (NED), as described by Chéry et al., (2014)^59^. For our analysis we defined a group “Other” which included samples in ADV_2 or NED. The GEO2R tool was used to find differentially expressed genes between ADV_1 and Other. A total of 56 probes (43 genes) with FDR < 0.05 were identified. These DEGs were further analyzed using Revigo^90^ to obtain functional gene ontology categories.

### Untargeted metabolomics

Metabolomic samples consisted of tumor cells collected from monoculture or co-culture with pericytes for 20 min. 1×10^6^ tumor cells were aliquoted and stored as a dry pellet at -80°C for metabolomics analysis. During expansion and collection of these samples, all cell culture materials were limited to a single batch to avoid technical differences that may be introduced by different media batches.

Untargeted metabolomics was performed by the Trans-NIH Metabolomics More. Ultra-high performance liquid chromatography (UHPLC) using the Bruker Elute Ultra-High-Performance Liquid Chromatograph was used to separate the chemicals prior to mass spectrometric analysis. Tandem mass spectrometry (MS/MS) and high-resolution mass spectrometry (HRMS) were performed using the Bruker tipstaff Pro Mass Spectrometer to determine structural information, molecular formulae and isotype patterns. Samples were analyzed using an ultra-high performance liquid chromatograph (Vanquish, Thermo Scientific) coupled to a high-resolution mass spectrometer (Orbitrap Fusion Tribrid, Thermo Scientific). LC-MS and LC-MS/MS data were acquired. LC-MS data were collected from individual samples (n = 1 injection), system blanks (injection of solvent used to resolubilize samples), and a pooled quality control. The pooled quality control (QC) was injected multiple times at different volumes and used in data processing. LC-MS/MS data, used to annotate features, were collected using the AcquireX (Thermo Scientific) deep scan methodology in which pooled QC was injected multiple times (n = 7). Ionization was performed via heated electrospray ionization (NG Ion Max, Thermo Scientific); data were collected in both positive and negative ionization modes. Data was obtained from 5 biological replicates each in 4 experimental groups.

Compound Discoverer 3.3.0.550 (Thermo Scientific) was used to process .raw files which resulted in a tabular output which included descriptors of each feature (e.g. m/z, retention time), annotation information (e.g. MS/MS database match), and peak area. We processed the output from Compound Discoverer using in-house R scripts via JupyterNotebooks. The major components of the processing included formatting of the data outputs, comparison of m/z and retention time of annotation features versus an in-house generated list based on authentic chemical standards, assessment of signal response in pooled QC samples, assessment of signal variance in pooled QC samples versus samples (i.e. dispersion ratio), and multi- and univariate statistics. MSI levels of annotation confidence were provided based on the MS/MS database matching algorithm in Compound Discoverer, a list of m/z generated from authentic chemical standards, and manual annotation. Features with an MS/MS database match were putatively annotated in alignment with the Metabolomics Standards Initiative level 2 annotations^91^. The following public, commercial, and in-house MS/MS spectral libraries were used to annotate features: NIST2020, GNPS, mzCloud, and in-house MS/MS spectral library acquired from authentic chemical standards purchased and run by the Metabolomics Core Facility.

Multivariate analysis was performed to identify relationships between experimental groups. Univariate analysis was used to analyze features independently, t-tests were performed on all features and groups of samples and multiple hypothesis testing correction was performed. Statistical significance was based on p-value <0.05 and log2 fold change of >1.

### Extracellular calcium depletion

Extracellular calcium depletion was achieved using low-calcium Hank’s solution (140 mM NaCl, 6 mM KCl, 5 mM D-Glucose, 1.3 mM MgCl_2_, 10mM Hepes in DI H_2_O), and compared directly to standard Hank’s solution containing 2 mM CaCl_2_. Pericytes were seeded in GA medium and allowed to adhere overnight. The following day, pericytes were labelled with 10 µM DiO and GA medium was replaced with either low- or normal-calcium hanks. Hank’s solutions were replaced every 15 min for 1 h.

Tumor cells were trypsinized, counted and incubated in either low- or normal-calcium Hank’s solution for 1 h, with solution replacement every 15 min. After incubation, tumor cells were added to pericytes in their respective calcium conditions and co-cultured for 2 h, followed by imaging to evaluate DiO transfer from pericytes to tumor cells.

### Calcium imaging

Pericytes were incubated in BrainPhys imaging optimized medium (05796, StemCell Technologies) containing 5 µM Calbryte (20651, AAT Bioquest) and 0.04% Pluronic F-127 (P-3000MP, Thermo Fisher Scientific) for 30 min at 37°C, followed by 15 min at room temperature. Calcium imaging of pericyte-tumor cell co-culture was performed using a Nikon SoRa spinning disk confocal microscope, acquiring images every 10 sec for a total duration of 10 min. To control for the pericyte calcium response to tumor-secreted factors, conditioned medium from 4T1 cells was applied to pericytes. Medium was conditioned by 4T1 cells for 20 min to match the co-culture time frame. Due to the asynchronous calcium signaling in co-culture, analysis was performed using the maximum change in fluorescence intensity per cell over the 10-minute recording period. Heatmaps were generated by first calculating Fo by averaging five of the lowest spatially binned ROI values, then subtracting Fo from F and dividing by Fo for each ROI. Heatmaps were produced using the imagesc function in Matlab.

Calcium imaging of pericytes in response to the Piezo1 agonist, Yoda1 (SML1558, Millipore Sigma), was performed using 5 min recordings at a rate of one frame every 3 sec. A 1 min baseline recording was acquired before the addition of either 30 µM Yoda1 or DMSO vehicle control, followed by a 3 min imaging. Finally, 10 µM ionomycin (I0634, Millipore Sigma) was added for 1 min as a positive control. Recordings from Yoda1 treated pericytes were analyzed using 10 cells per image and each time point was background corrected using the average of 10 background ROIs at the same time point. Calcium flux values were normalized to T0 fluorescence intensity.

### Mechanostimulation induced phospholipid release

Pericytes were loaded with DiO in GA media for 1 h at 37°C. Following labeling, pericytes were treated with 30 µM Yoda1, DMSO vehicle, or left untreated for 20 min. Conditioned medium (CM) from pericytes was collected post-treatment. Tumor cells were then incubated with the pericyte CM for 20 min, after which the medium was removed by centrifugation to simulate a 20 min co-culture exposure. These cells were then seeded onto plastic for 48 h to assess proliferation. For lipid transfer analysis, an additional set of tumor cells was incubated with pericyte CM for 2 h to evaluate the transfer of DiO-labeled phospholipids derived from pericytes following pharmacological activation of Piezo1 channels.

### Drug screen to inhibit lipid transfer

Compounds screened for ability to inhibit phospholipid transfer included nitric oxide synthase 2 (Nos2) inhibitor Aminoguanidine (0.5 mM), cyclooxygenase 2 (Cox2) inhibitor Indomethacin (50 mM), and calcium channel inhibitors, Gadolinium (5 mM), Thapsigargin (3 nM) and Nimodipine (100 mM). Pericytes were seeded on a 96-well black clear bottom plate in GA medium to adhere overnight. The following day, pericytes were labeled with 10 µM DiO and pre-treated with the selected compound or vehicle control for 1 h prior to co-culture.

Tumor cells were trypsinized, counted and pre-treated with the selected drug or vehicle for 1 h. Tumor cells were added to pericytes at a 1:1 ratio and co-cultured for 2 h. Imaging was performed to assess DiO transfer from pericytes to tumor cells.

### Bioorthogonal (Click) chemistry

To assess the transfer of pericyte derived proteins, pericyte growth medium was supplemented with Click-iT Homoproparglyglycine (HPG) reagent (C10428, Thermo Fisher Scientific) for 5 days to allow incorporation of analogue during protein synthesis. After expansion with or without HPG, pericytes were seeded in a 96-well plate and allowed to adhere overnight in GA medium.

Tumor cells were trypsinized, counted and seeded in monoculture or co-culture with the pericytes for 2 h. A portion of tumor cells were plated in the presence of HPG for 30 min as a positive control. After 2 h the medium was removed, and the cells were fixed with 3.7% paraformaldehyde (PI28906, Fisher Scientific). The cells were blocked, permeabilized and incubated with the click reaction cocktail (C10428, Thermo Fisher Scientific) for 30 min at room temperature, protected from light. Finally, nuclei were stained with a nuclear mask before imaging.

### Data Analysis

Data are presented as mean ± standard deviation (SD). Where relevant, statistics were performed on the mean values of the independent experiments. Statistical tests were performed using GraphPad Prism 5 software and statistical significance was accepted at the 95% confidence level. Image analysis was performed using Gen5, ImageJ or Halo.

## Data Availability

RNA-Seq data that support the findings of this study will be deposited in the Gene Expression Omnibus (GEO). Publicly available RNA-Seq of DTC in bone of prostate cancer patients were used and can be found in ref^59^ and under GEO accession number GSE48995.

## Acknowledgements

The NCI CCR genomics core and NCI sequencing facility performed RNA sample quantification, quality measurements and sequencing. The Trans-NIH Metabolomics core performed the mass spectrometry experiment and annotation of metabolites. Histopathology laboratory at the National Laboratory for cancer research performed cryosectioning. The CCR microscopy core provided training on microscopes and troubleshooting assistance. Devorah Gallardo performed intracardiac injections. Wendy Dubois performed animal genotyping and colony maintenance. Nima Ghitani generated heat maps of calcium flux analysis.

## Author Contributions

T.M., M.J.G., E.C.M., C.A.M., and L.E.Z. conducted the experiments. M.J.K. assisted with confocal microscopy experiments. A.K.J. and K.E.O. performed mass spectrometry and metabolite annotations. M.P.L. and H.H.Y. generated DEG lists from publicly available datasets. A.T.C. provided expertise regarding PIEZO1 biology. T.M. and M.M. prepared to manuscript draft. M.M. and C.M.B. designed and supervised the project. All authors reviewed and approved the final version of the manuscript.

