## Supplementary figures and images for "Pericyte-tumor crosstalk facilitates metastatic tumor cell latency through PIEZO1-activated lysophospholipid transfer"

### Supplemental Figures 1-4

#### Supplemental Figure 1

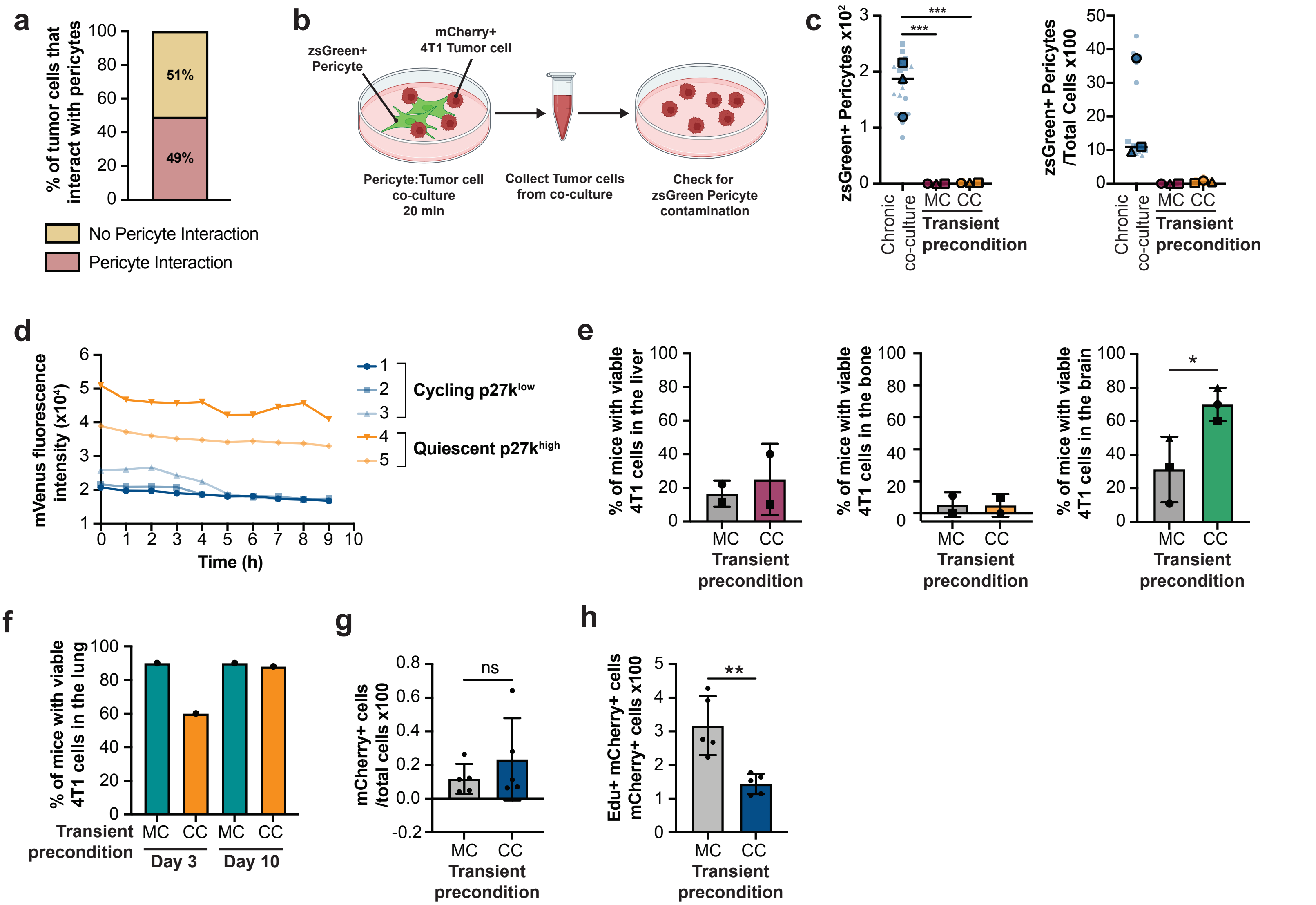

#### Supplemental Figure 2

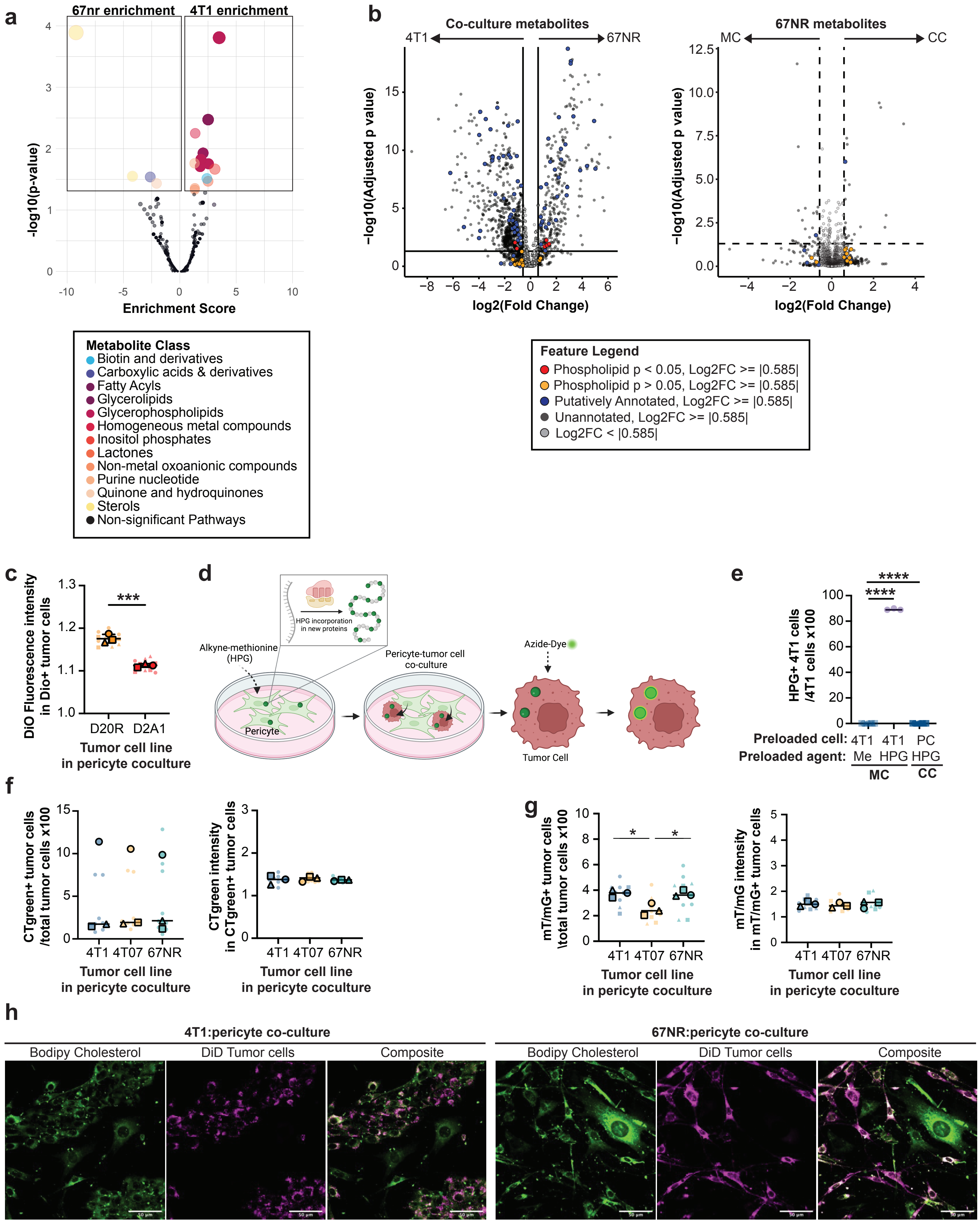

#### Supplemental Figure 3

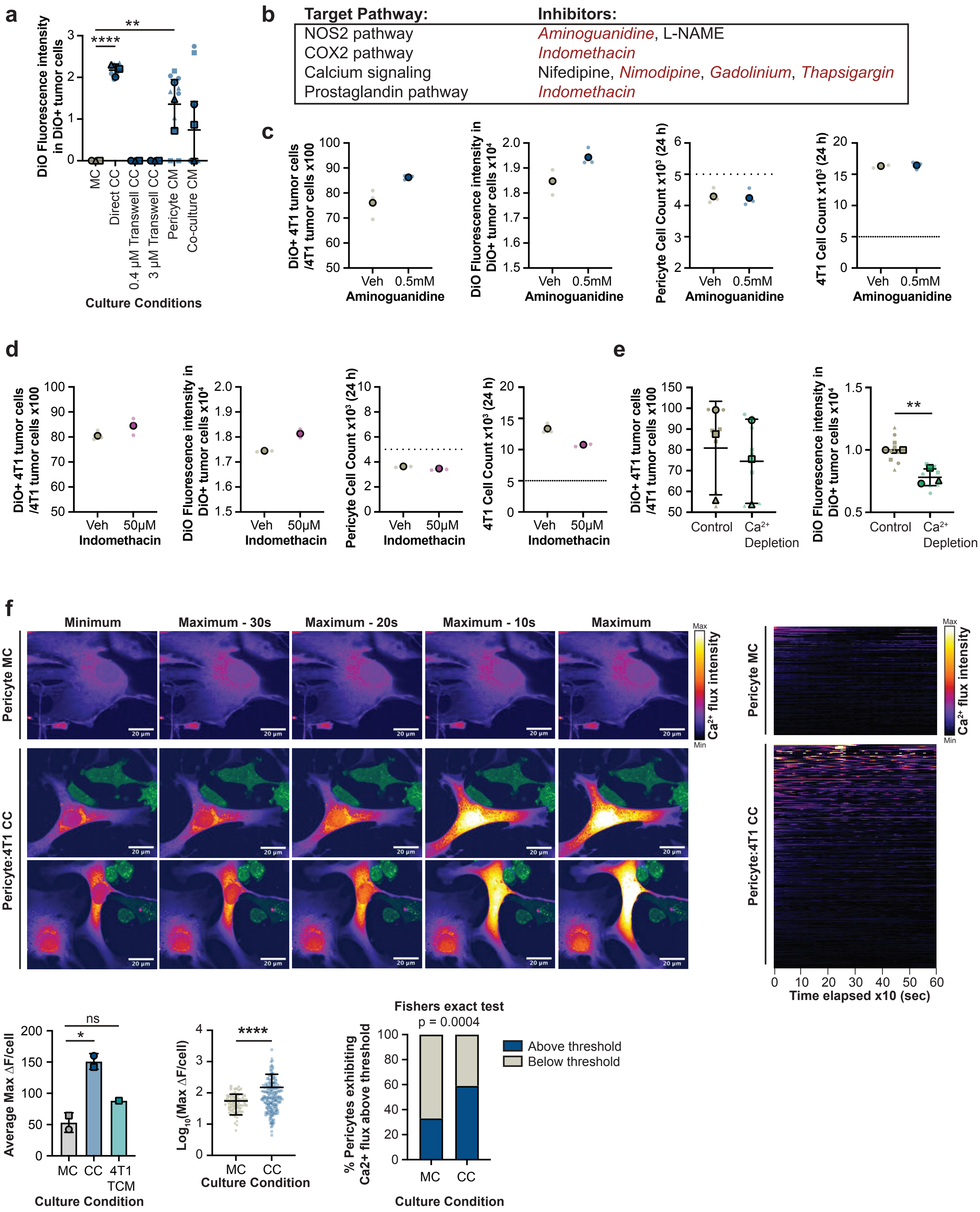

#### Supplemental Figure 4

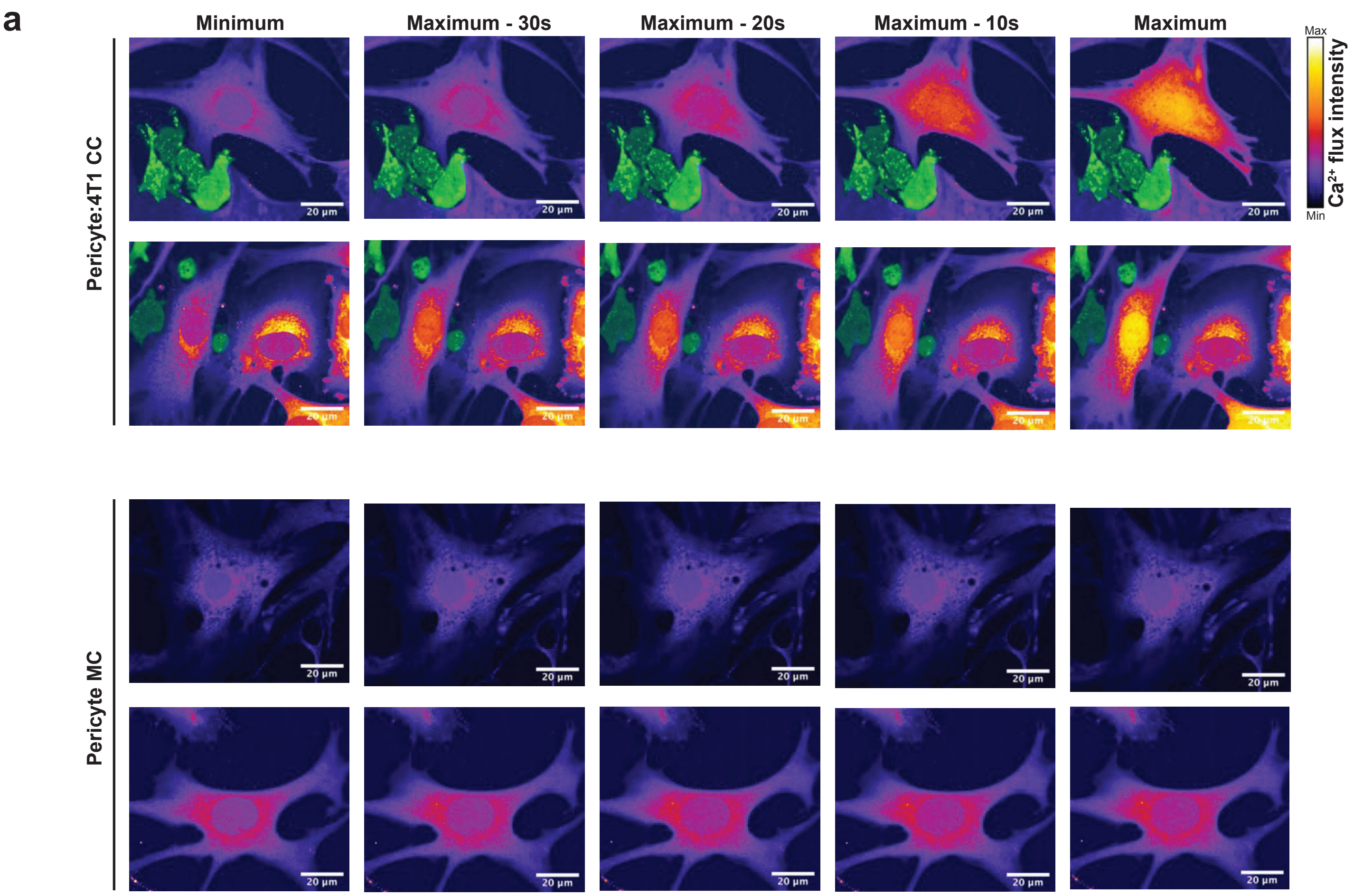


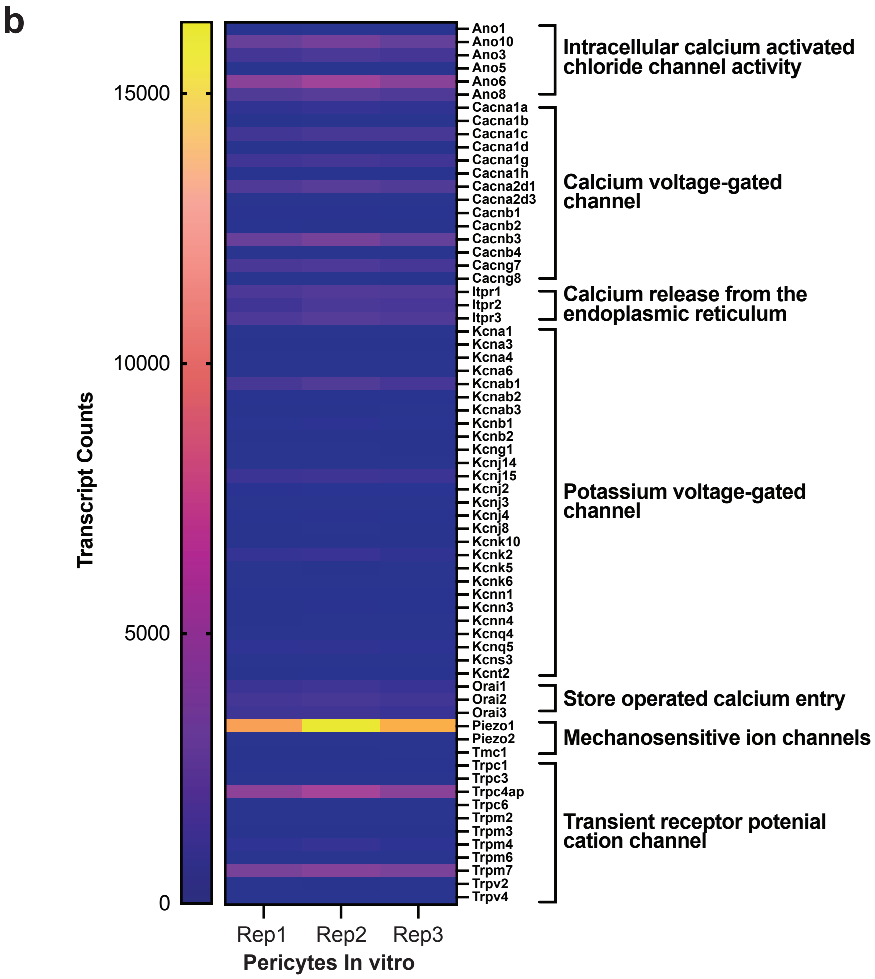
